# p63 sets the threshold for induction of apoptosis using a kinetically encoded ‘doorbell-like’ mechanism

**DOI:** 10.1101/681007

**Authors:** Jakob Gebel, Marcel Tuppi, Apirat Chaikuad, Katharina Hötte, Laura Schulz, Frank Löhr, Niklas Gutfreund, Franziska Finke, Martin Schröder, Erik Henrich, Julija Mezhyrova, Ralf Lehnert, Francesco Pampaloni, Gerhard Hummer, Ernst H.K. Stelzer, Stefan Knapp, Volker Dötsch

**Author notes:** These authors contributed equally to this work (Jakob Gebel, Marcel Tuppi, Apirat Chaikuad). Corresponding authors. Institute of Biophysical Chemistry, Centre for Biomolecular Magnetic Resonance, University of Frankfurt, Max-von-Laue-Strasse 9, 60438 Frankfurt/Main, Germany.; or The Francis Crick Institute, London, NW11ST, UK. **Abbreviations**: DSBs, DNA double strand breaks; POI, premature ovarian insufficiency; TAD, transactivation domain; DBD, DNA binding domain; TD, tetramerization domain; SAM, sterile alpha motif; PAD, phosphorylation activation domain; TID, transcriptional inhibitory domain; NMR, nuclear magnetic resonance; Dox, doxorubicin; CHK2, checkpoint kinase 2; CK1, casein kinase 1; Cs, Cisplatin; PARP 1, poly(ADP-ribose)-polymerase; DSBs, double strand breaks.

## Abstract

Cell fate decisions such as apoptosis require cells to translate signaling input into a binary yes/no response. A tight control of the process is required to avoid loss of cells by accidental activation of cell death pathways. One particularly critical situation exists in primary oocytes because their finite number determines the reproductive capacity of females. On the one hand a stringent genetic quality control is necessary to maintain the genetic integrity of the entire species; on the other hand an overly stringent mechanism that kills oocytes with even minor DNA damage can deplete the whole primary oocyte pool leading to infertility. The p53 homolog TAp63α is the key regulator of genome integrity in oocytes. After DNA damage TAp63α is activated by multistep phosphorylation involving multiple phosphorylation events by the kinase CK1, which triggers the transition from a dimeric and inactive conformation to an open and active tetramer. By measuring activation kinetics in ovaries and single site phosphorylation kinetics *in vitro* with peptides and full length protein we show that TAp63α phosphorylation follows a biphasic behavior. While the first two CK1 phosphorylation events are fast, the third one that constitutes the decisive step to form the active conformation is slow. We reveal the structural mechanism for the difference in the kinetic behavior based on an unusual CK1/TAp63α substrate interaction and demonstrate by quantitative simulation that the slow phosphorylation phase determines the threshold of DNA damage required for induction of apoptosis.

## INTRODUCTION

The reproductive lifespan of women is determined by the primordial follicle (PF) reserve. PFs consist of primary oocytes surrounded by a single flat layer of granulosa cells. These oocytes are arrested in prophase I of meiosis I until being recruited for ovulation. Menopause is initiated in humans when the number of PFs decreases from its original level of one to two million at the time of birth to below 1000^1^. Depletion of the PF reserve was identified as the major cause for premature ovarian insufficiency (POI). In female patients suffering from cancer, sickle cell anemia or certain autoimmune diseases, treatment with chemotherapeutic drugs and/or irradiation can deplete the ovarian reserve, resulting in POI^2,3^. Fewer than ten DNA double strand breaks (DSBs) (γ-irradiation with 0.45 Gy) are suffice to eliminate the entire PF reserve in mice^4^. In humans, the LD_50_ total body irradiation dosage for the loss of the PF reserve was extrapolated to be less than 2 Gy, while the typical total body irradiation dose for acute leukemia patients is 12 Gy^5,6^. The loss of the PF reserve also causes the breakdown of the endocrine function of the ovary resulting in addition to infertility in health impairments like osteoporosis, cardiovascular disease and psychosocial disorders in addition to infertility^7^. In contrast to the arrested PFs, growing follicles are unaffected from low dosage of irradiation. The decisive difference between both oocyte types is the expression of TAp63α, a p53 orthologue, in PFs^4,8^. The expression of TAp63α starts shortly after the pachytene stage of meiosis I while oocytes that have reentered the cell cycle are devoid of TAp63α^4^.

The high levels of the pro-apoptotic factor TAp63α combined with the long arrest time of oocytes (up to 50 years in humans) require a very tight regulation of TAp63α’s activity to avoid cell death of uncompromised oocytes. In contrast to all other members of the p53 family that form tetramers through an oligomerization domain, TAp63α adopts a closed, inactive and only dimeric conformation^9^. DNA damage triggers a kinase cascade resulting in the phosphorylation of TAp63α, which disrupts the autoinhibitory dimeric complex and triggers the formation of an open active and tetrameric conformation resulting in the elimination of the damaged oocyte^10^. Recently, we and others have identified the kinases involved in this process, mapped the phosphorylation sites and described the structural mechanism of the activation process. The first step of the kinase cascade is the activation of checkpoint kinase 2 (CHK2) or checkpoint kinase 1 (CHK1) by ataxia telangiectasia mutant kinase (ATM)^11^. Activated CHK2 phosphorylates TAp63α on S582^12^ which renders TAp63α a substrate for casein kinase 1 (CK1)^13^. CK1 requires pre-phosphorylated substrates with the consensus sequence pS/T-x-x-S/T, where pS/T is a phosphorylated serine or threonine. In TAp63α CK1 adds four phosphate groups that lead to the disruption of the autoinhibitory complex through electrostatic repulsion^13^. Our previous experiments have shown that the inhibited dimeric state of TAp63α constitutes a kinetically trapped high energy state^14^. Consequently, activation follows a spring-loaded mechanism that explains the high sensitivity of oocytes towards DNA damage. This activation mechanism has to be adjusted to a certain level of damage that on the one hand must be sufficiently low to protect the integrity of the genetic pool of a species but on the other hand tolerant enough not to endanger reproductive capacity. This situation requires a mechanism that ideally works similar to a doorbell: an input signal (mechanical pressure) below a certain threshold has no effect. If, however, this threshold is surpassed the output signal is independent of the actual input signal strength (pressing harder does not make the doorbell louder). Indeed, a tight dose-response curve has been measured in four-day old mice: while most oocytes survive irradiation with 0.1 Gy (~three DSBs per cell), virtually all primary oocytes were eliminated by 0.45 Gy irradiation (~ten DSBs per cell)^4^.

Switch-like processes are often based on the integration of two different and independent signals. An accidental activation is thus suppressed as the likelihood of activation is the product of two small probabilities. Examples include regulation of actin polymerization via N-WASP which requires co-stimulation by Cdc42 and PIP2^15^ as well as activation of T-cells via triggering nuclear import of NFAT and activation of the transcription factor AP1^16^. The activation of TAp63α is based on phosphorylation by two kinases which in principle could provide such a sigmoidal activation. However, both act in a sequentially dependent manner and CK1 kinases are thought to be constitutively active^17^. These properties would prevent a switch-like activation and make the initiation of apoptosis dependent only on CHK2 with potentially detrimental consequences for the safety of oocytes. To understand how TAp63α converts a graded stress response into a doorbell like activation mechanism we investigated the kinetics of apoptosis in primary oocytes both at the cellular and molecular level.

## RESULTS

### DNA damage induced apoptosis in primordial follicles follows a sigmoidal time response

We set out to measure the time dependent TAp63α activation in whole mouse ovaries following DNA damage as a response to γ-irradiation (0.5 Gy) as a direct source of DSB induction. The tetramerization kinetics of TAp63α upon γ-irradiation followed a sigmoidal time response with virtually all TAp63α converted into its active tetrameric state within two hours (Fig. 1a, 1b and Supplementary Fig. 1a). Activation can be suppressed using selective inhibitors of ATM, CHK2 and CK1, suggesting that tetramerization is based on the same pathway as activation following the treatment with cisplatin or doxorubicin (Supplementary Fig. 1a)^13,14,18^. In order to investigate the time dependent induction of apoptosis, we measured the ratio of germ cell nuclear acidic peptidase (GCNA)^19^ positive and cleaved poly(ADP-ribose)-polymerase 1 (PARP1) double positive cells in whole mouse ovaries using our 3D staining and light sheet-based fluorescence microscopy combined with semi-automated segmentation method (Fig. 1c and Supplementary Fig. 1b)^13^. The primary oocyte specific GCNA^10^ staining allows the specific quantification of apoptosis markers in primary oocytes. Cleaved PARP1 was detected starting from four hours after irradiation and reached 100% in the remaining PFs at ten hours showing a sigmoidal overall transition (Fig. 1d). A sigmoidal transition was also evident when monitoring the decline of the integrated GCNA signal in whole ovaries representing the total number of remaining primary oocytes (Fig. 1e).

**Fig. 1.**
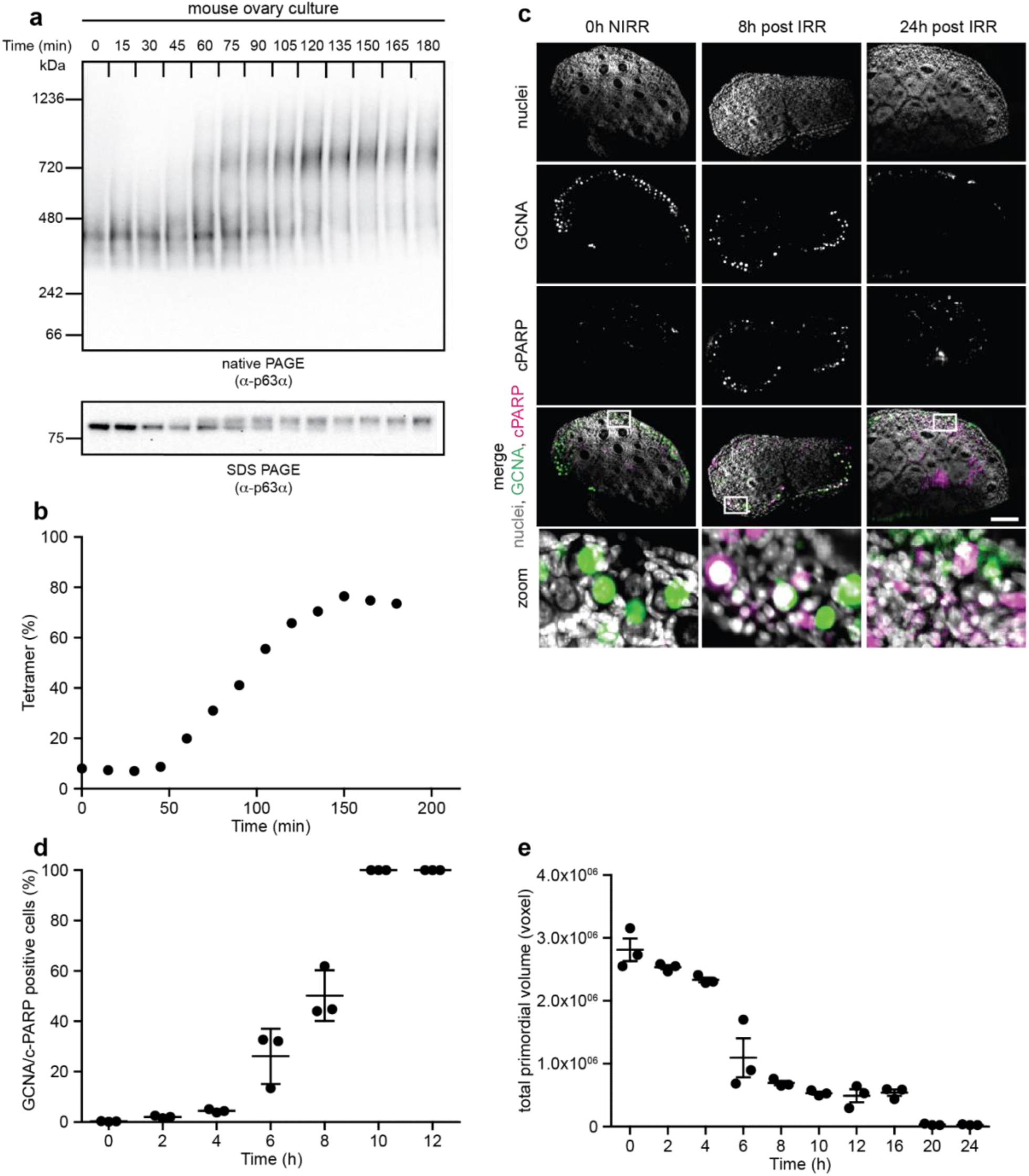
TAp63α mediated primordial follicle death follows an overall sigmoidal kinetics. **a**, Whole mouse ovaries were *ex vivo* γ-irradiated with 0.5 Gy or left untreated. The oligomeric state was analyzed as a function of time by BN-PAGE and the phosphorylation dependent mobility shift by SDS-PAGE. **b**, Densiometric analysis of the dimeric and tetrameric fraction of **a. c**, Representative single plane images of whole mount 3D ovary stainings using DAPI (nuclei), GCNA and cleaved PARP (c-PARP) at the indicated time points after the 0.5 Gy γ-irradiation (IRR) and of untreated ovaries. **d**, Ratio of c-PARP positive GCNA positive cells of whole ovaries after 0.5 Gy γ-irradiation using whole mount 3D ovary staining. **e**, Time dependent GCNA positive cells, quantified in voxels, after 0.5 Gy γ-irradiation using whole mount 3D ovary staining. **d** and **e**, data presented as mean ± SD (*n* = 3). See also Supplementary Fig. 1.

### Phosphorylation of the third CK1 site is the slowest and the ‘point of no return’

The correlation between the sigmoidal time response patterns for TAp63α tetramerization and induction of apoptosis suggests that the molecular activation mechanism of TAp63α is the decisive process controlling the fate of the oocyte. Earlier, we have shown that the initial phosphorylation of S582 by CHK2 is necessary but does not result in active TAp63α. This phosphorylation serves as the priming event for recruiting CK1 which adds phosphate groups at four consecutive sites: S585, S588, S591 and T594^13^. We also have characterized the importance of each phosphate group for the activation mechanism by mutating the individual residues that can be phosphorylated to alanine and measuring the tetramerization kinetics following addition of CK1. Demonstrating that the last phosphorylation event, T594, is dispensable for TAp63α activation. However, mutating S591 abrogates the conversion of TAp63α to a tetramer, showing that phosphorylation of S591 is ‘the point of no return’ for TAp63α activation^13^. This observation raised the question how phosphorylation of the three CK1 sites, S585, S588 and S591 can provide a switch-like activation mechanism. Detailed theoretical and experimental studies have shown that multisite phosphorylation can lead to ultrasensitivity^20–22^. This is achieved either by having strongly differing kinetic constants for the individual reaction steps or if the process is distributive^23^. In addition, a distributive mode of action would allow the interference of a phosphatase while a processive mode of action would result in a continuous and undisturbed phosphorylation of all sites. To investigate the mode of action we analyzed the kinetics of phosphorylation of the individual sites within a peptide, that corresponds to the sequence N-terminal to the transactivation inhibitory domain (TID)^24^ and includes the phosphorylation sites for CHK2 and CK1 (hereafter referred to as phosphorylation activation domain (PAD)) via nuclear magnetic resonance (NMR) spectroscopy. Phosphorylation of S582 by MapKap kinase 2 (MK2) which recognizes the same phosphorylation sequence as CHK2 resulted in the appearance of a new resonance in the HSQC spectrum and loss of the original S582 signal, consistent with the peptide being phosphorylated on this serine residue (Supplementary Fig. 2a). MK2 can be easily produced in bacteria, is constitutively active and therefore a convenient surrogate for CHK2 *in vitro* priming of substrates.

Recording the phosphorylation kinetics of this pre-phosphorylated peptide by CK1δ showed a biphasic behavior. Phosphorylation of S585 and S588 occurred very quickly with an almost indistinguishable kinetics, which corresponds to either a processive mode or to a distributive mode with a very fast product releasing and substrate re-binding kinetics. A stark contrast was observed for the subsequent phosphorylation of S591 with a ~40–fold slower rate (Fig. 2a, 2B and Supplementary Table 1). The observed increase in the concentration of the triple phosphorylated peptide (pS582, pS585, pS588; PAD-3P) beyond the concentration of the kinase implies that CK1 must have dissociated from the PAD-3P product peptide to phosphorylate S585 and S588 in free PAD-1P and PAD-2P peptides. Only when the supply of pS582 mono-phosphorylated peptide is exhausted, S591 and T594 become substrates for CK1. We conclude from these observations that 1) phosphorylation of the entire stretch occurs via a distributive mode and that 2) S585 and S588 act as a buffer that delays phosphorylation of the downstream sites S591 and T594. In the same time frame as the phosphorylation of S591 and T594 a non-consensus sequence phosphorylation of T586 appears. The T586A mutant showed no difference in the phosphorylation kinetics demonstrating that phosphorylation of T586 is not the reason for the biphasic behavior (Supplementary Fig. 2h). Interestingly, phosphorylation of S591 is the point of no return for the activation and the kinetic experiment demonstrate that this critical phosphorylation is the slow step occurring in a distributive mode of action.

**Fig. 2.**
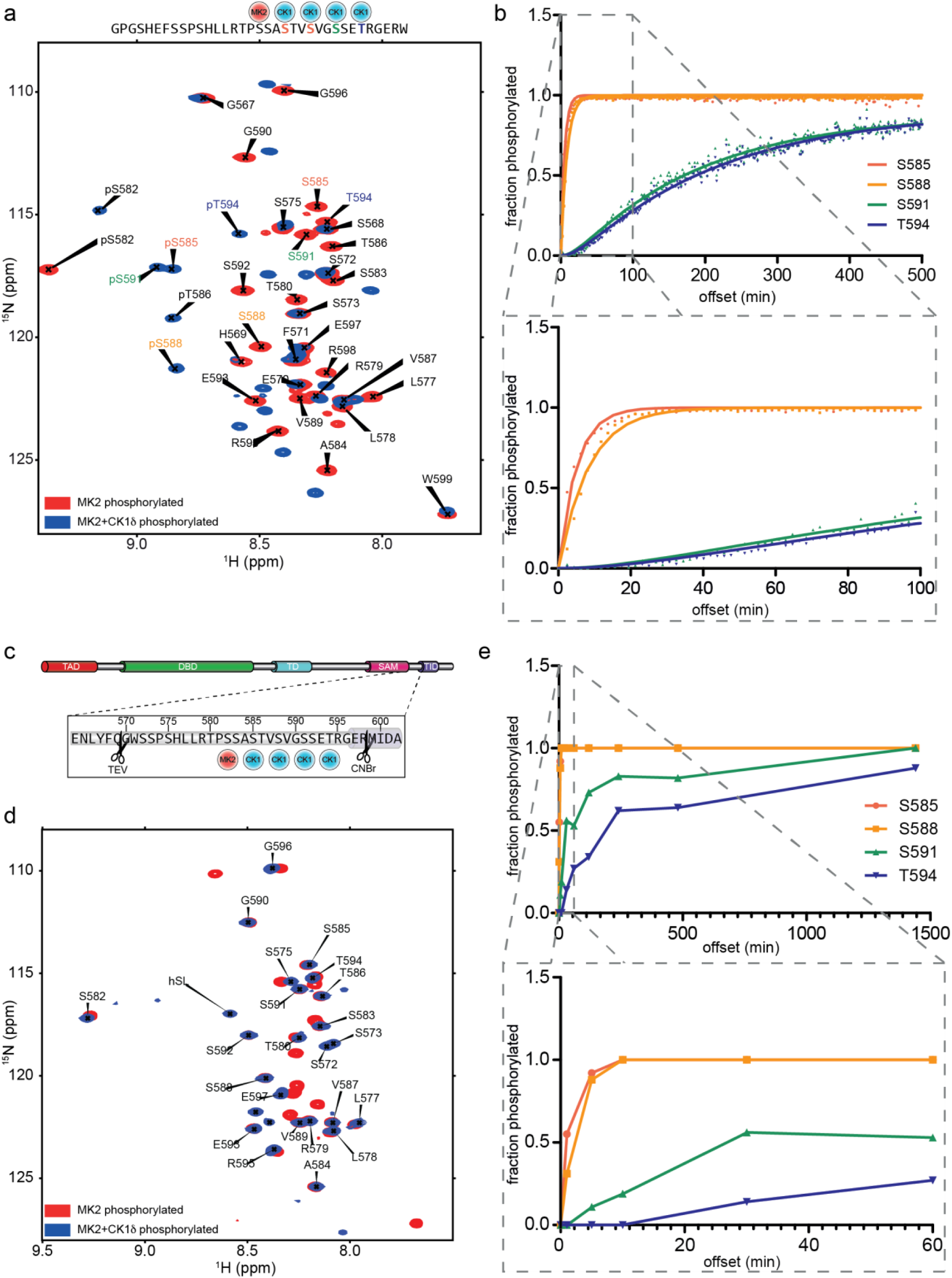
The third CK1 phosphorylation is the slowest step in the TAp63α phosphorylation and constitutes the ‘point of no return’. **a**, Single site phosphorylation kinetics was measured by NMR spectroscopy. Overlay of the [^15^N, ^1^H]-HSQC spectra of MK2 pre-phosphorylated (red) and CK1 phosphorylated (blue) spectra of the PAD peptide representing the starting and the end point. The analyzed peptide sequence is shown above the spectra. **b**, Quantitative evaluation of the phosphorylation kinetics of the four CK1 sites. Zoomed graph at the bottom focuses on the initial reaction showing that the third and fourth sites get modified only after the first two sites are almost 100% phosphorylated. **c**, Schematic representation of the domain structure of TAp63α showing the location of the PAD peptide and the sites used for cleaving the peptide by TEV protease and CNBr. **d**, Overlay of [^15^N, ^1^H]-HSQC spectra of MK2 pre-phosphorylated (red) and CK1 phosphorylated (blue) PAD peptide cleaved from full-length TAp63α. **e**, Quantitative evaluation of the phosphorylation kinetics of the four CK1 sites in full-length TAp63α. See also Supplementary Fig. 2 and Supplementary Table 1.

### The phosphorylation kinetics in the full-length protein mirrors the kinetics of the isolated peptides

Phosphorylation kinetics of the isolated PAD peptide might, however, be quite different from the kinetics of the corresponding peptide within the closed dimeric full-length TAp63α conformation since the sequence is surrounded by secondary structure elements and folded domains (Fig. 2c). We measured the phosphorylation kinetics with ^15^N-labeled full-length dimeric TAp63α. To be able to analyze the phosphorylation kinetics by NMR we stopped the reaction by adding a high concentration of EDTA and CK1 inhibitor PF-670462. Subsequently, we isolated the PAD peptide first by cleaving with tobacco etch virus (TEV) protease and after isolating the resulting C-terminal fragment the PAD was cleaved from the TID using CNBr. For this purpose, a protease cleavage site was engineered N-terminal to the PAD and a V599M mutation at its C-terminal end (Supplementary Fig. 2b-g). The phosphorylation kinetics of the PAD peptide and the PAD derived from the dimeric TAp63α showed a similar pattern. S585 and S588 were phosphorylated fast, while phosphorylation of S591 and T594 were significantly slower (at least 25-fold) following a distributive mechanism (Fig. 2d and 2e). This demonstrated that the biphasic activation mode also occurs in the full-length dimeric TAp63α and is not a peptide derived artefact.

### The biphasic kinetics is p63 sequence specific

The observed biphasic behavior could be a property of the p63 sequence, of CK1 substrate recognition or a mixture of both properties. Kinases of the CK1 family recognize several hundred confirmed or suggested substrates. Among them are the period proteins (PER1-3), β-catenin, NF-AT1,2,4, adenomatous polyposis coli (APC), p53, MDM2 and Yes-associated protein 1 (YAP1)^17^. Some of these targets contain stretches of phosphorylatable residues harboring the CK1 consensus target sequence and these sequences are known to get phosphorylated at multiple sites (Fig. 3a). To investigate if CK1δ can in principle add more than two phosphate groups in a processive manner we chose YAP1 which contains a phosphodegron sequence that regulates its cellular localization and degradation. YAP1 is pre-phosphorylated by LATS kinase at S397, within the HxRxxS consensus motif. CK1 gets recruited to phosphorylated YAP1 and adds three more phosphate groups in the sequence HSRDESTD**S**GL**S**MS**S**YS^25^. We used a modified, MK2 phosphorylatable peptide and measured the NMR-based kinetics of the three CK1 phosphorylation events after MK2 priming (Fig. 3b, 3c, Supplementary Fig. 3a and 3b). In a stark contrast to the PAD peptide, the results demonstrated fast kinetics for all three CK1 sites, suggesting that CK1 phosphorylates the sequence in YAP1, and possibly other proteins, without a distinct biphasic mechanism (Fig. 3d, 3e and Supplementary Table 1).

**Fig. 3.**
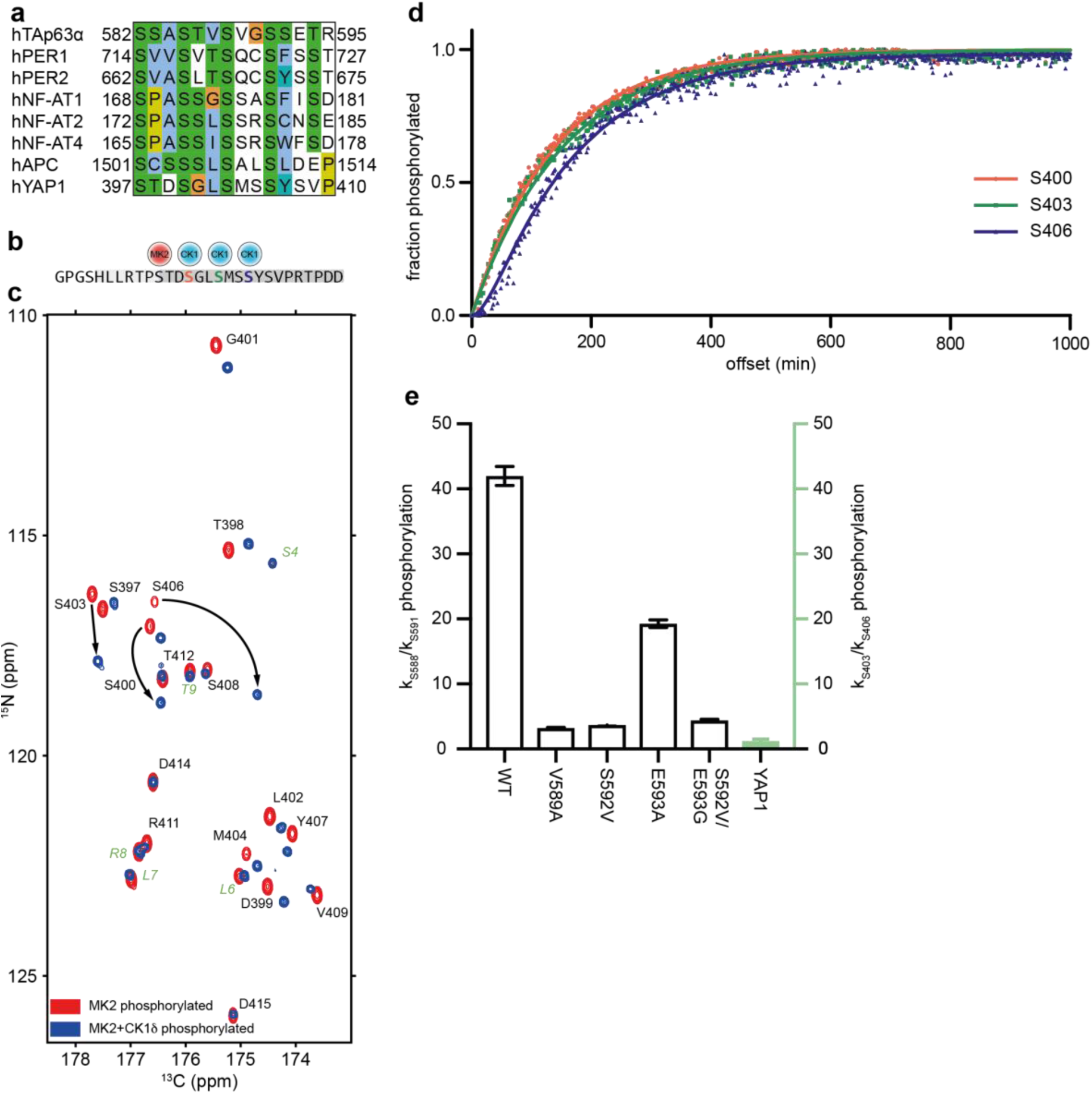
Kinetics of YAP1 phosphorylation by CK1 is not biphasic. **a**, Comparison of sequences taken from the indicated proteins that harbor multiple CK1 phosphorylation sites. Sequences refer to hTAp63α: NM_003722.4 (numbering according to AF_075430.1), hPER1: NM-002616.2, hPER2: NM_022817.2, hNF-AT1: NM_173091.3, hNF-AT2: NM_001278669.1, hNF-AT4: NM_173165.2, hAPC: NM_000038.6 and hYAP1: NM_001130145.2. **b**, Sequence of the YAP1 peptide highlighting the site of pre-phosphorylation (red) and the three CK1 sites (blue). **c**, Overlay of 2D ^13^C-detected (H)NCO spectra of MK2 pre-phosphorylated (red) and CK1 phosphorylated (blue) YAP1 peptide. Arrows show representative examples of signals shifted due to phosphorylation. Amino acids not belonging to the YAP1 sequence are indicated in green. **d**, Quantitative evaluation of the kinetics of the three CK1 sites demonstrating that the three phosphorylation sites show a very similar kinetics. **e**, Bar diagram describing the difference in rate constants of S588 and S591 phosphorylation in wild type p63 and mutants as well as YAP1 (green bar, right side). The error represents the standard deviation of the fitting procedure. See also Supplementary Fig. 3 and Supplementary Table 1.

### The CK1 p63 PAD complex structure reveals the basis for the biphasic activation kinetics

This result suggested that the p63 PAD sequence is responsible for the observed biphasic kinetics and we hypothesized that the residues C-terminal to the third CK1 phosphorylation site (S591) might be crucial. We exchanged the sequence following the third CK1 site S591 (S592 and E593) with the sequence directly C-terminal to the second CK1 phosphorylation site S588 (V589 and G590). Interestingly, phosphorylation of S591 and T594 in this double mutant was indeed ten-fold accelerated in comparison to wild type. A faster phosphorylation kinetics was evident also in both single S592V and E593A mutants with a notably more pronounced effect observed for the S592V mutation (Fig. 3e, Supplementary Fig. 3c, 3d and Supplementary Table 1).

To obtain a mechanistic insight of the kinase–peptide interactions that are responsible for the delayed kinetics, we determined the crystal structures of the kinase domain of CK1δ in complexes with AMPPCP/ADP and various PAD peptides harboring different phosphorylation states, including PAD-1P (the first substrate with primed phosphorylation at pS582), PAD-2P (pS582 and pS585) and PAD-3P (pS582, pS585 and pS588) (Supplementary Fig. 4a-f). The overall conformations of all bound peptides were highly similar, yet they captured different states, including the substrate-bound form in PAD-1P and PAD-2P having the S585 and S588, respectively, located at the catalytic site. In contrast to PAD-1P/2P the PAD-3P crystalized in a product-PAD-3P complex in which pS588 was positioned within the catalytic site (Fig. 4a, 4b and 4d). Structural analyses of the CK1δ-PAD-2P complex as an example of the substrate-bound state revealed that the interactions between the kinase and the peptide were induced by electrostatic charge complementarity of both anchor-ends on the N- and C-termini of the peptide (Fig. 4c).

**Fig. 4.**
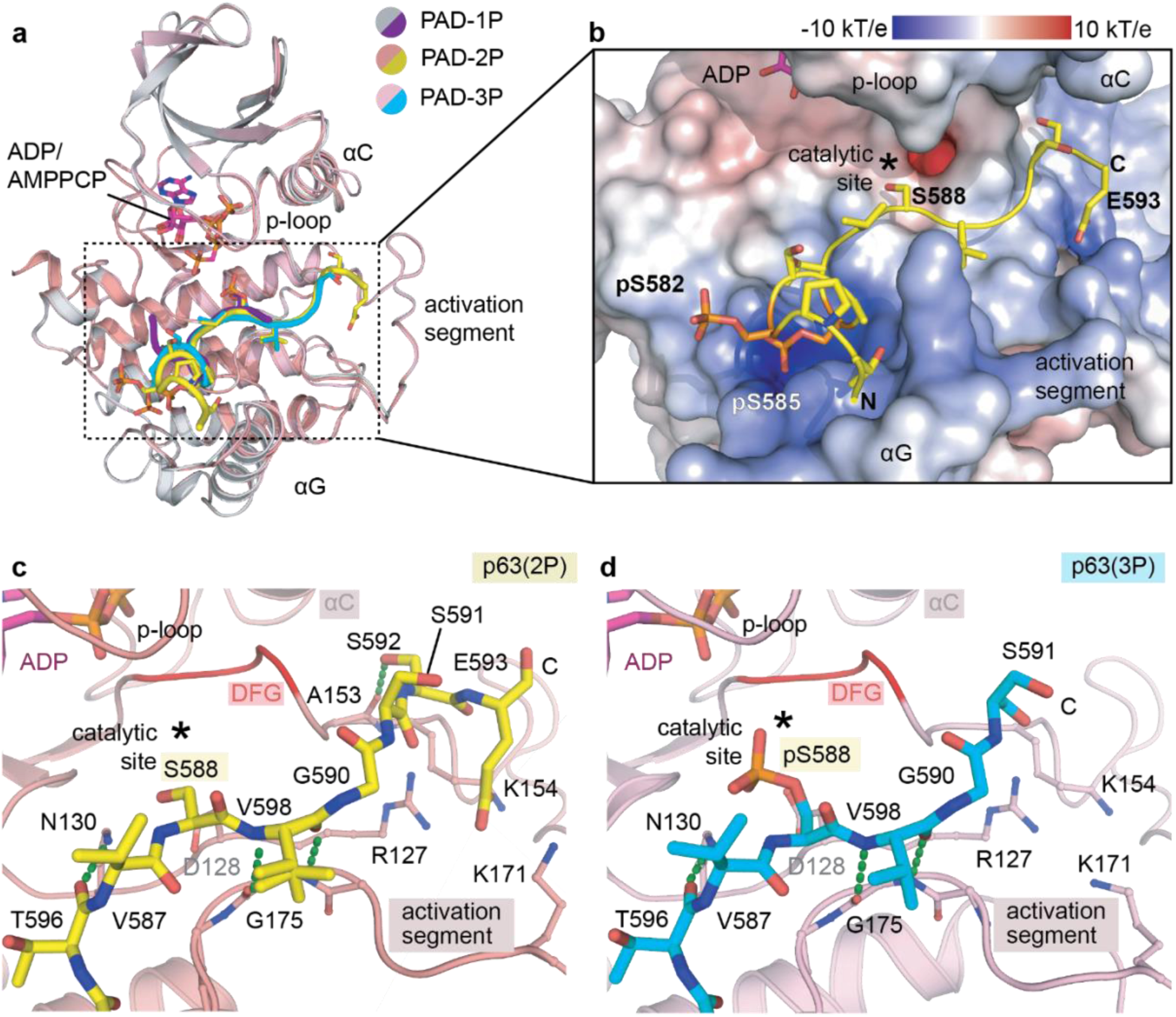
Crystal structures of CK1δ in complexes with different PAD peptides. **a**, Superimposition of all complexed structures demonstrates similar binding conformation of the PAD peptides within the kinase. Three PAD peptides colored differently as indicated harbor different phosphorylation states, including PAD-1P containing single phosphorylated S582, PAD-2P with double phosphorylations on S582 and S585, and PAD-3P with triple phosphorylations on S582, S588 and S588. **b**, Electrostatic potential on the kinase surface at the substrate peptide binding sites reveals both charge and shape complementarity for an accommodation of the PAD-2P peptide. **c-d**, Detailed interactions between the PAD-2P or PAD-3P and the kinase are highly resembled despite representing different states of substrate- and product-bound complexes, respectively. See also Supplementary Fig. 4 and Supplementary Table 2.

The peptide N-terminal part adopted a helical turn positioning the phosphate groups of pS582 and pS585 into the basic pocket of the kinase formed by Arg178 and Lys224 (for clear referencing, the single amino acid letter code is used for the PAD peptide and the three-letter code for the CK1δ kinase), while the PAD C-terminal E593 residue interacted with a second basic cluster present in the CK1δ substrate binding groove. This second cluster was formed by the N-terminus of the activation segment of the kinase. Furthermore, a potential hydrogen bond between the PAD S592 and the backbone carbonyl of CK1δ Ala153 was formed. The middle stretch of the PAD central to S588 at the catalytic site, displayed a canonical kinase-substrate interaction (Fig. 4c and 4d). Close inspection of the region surrounding the catalytic site revealed a non-polar patch on the kinase that was compatible with a medium-sized hydrophobic substrate residue such as V589 of the PAD peptide, which was tucked within a groove fenced on one side by the αG of the kinase. This explains the preference for hydrophobic amino acids in CK substrates at the +1-position relative to the serine/threonine phosphorylation site (Fig. 4c and 4d). In comparison, the binding mode of the PAD-3P in a product bound state highly resembles that of the substrate PAD-2P, albeit with the positioning of pS588 at the catalytic site with its phosphate moiety adjacent to the DFG Asp149 and the HRD Asp128 facilitated by an unusually bent ADP conformation (Fig. 4c and 4d). These structural insights from both substrate- and product-bound states suggested that such accommodation of the PAD peptides with S588/pS588 at the catalytic sites likely offered an optimal binding mode within the kinase, presenting the charge compatibility not only at N- and C-terminal anchor-points but also the non-polar groove for the middle part.

Processivity towards the next phosphorylation event on S591 requires the movement of the triple phosphorylated peptide by a translocation of S592 and E593 away from their favorable binding polar/charged pocket to the unfavorable hydrophobic environment, explaining the accelerated kinetics of the mutants (Fig. 3e). To further investigate this model, we measured the phosphorylation kinetics for the V589A mutant which replaces a large hydrophobic residue with a smaller one. This mutation indeed slowed phosphorylation of the second phosphorylation site ~three-fold probably by weakening the interaction with the small hydrophobic surface in the CK binding site. At the same time, the mutant V589A also accelerated phosphorylation of S591 approximately threefold (Fig. 3e, Supplementary Fig. 3c, 3d and Supplementary Table 1). Replacing the medium sized hydrophobic valine residue with a smaller alanine probably weakened the interaction in this +1-substrate position, resulting in both suboptimal positioning and locking of the substrate and a smaller hurdle for the release of the product.

We further investigated the interaction between the kinase and the PAD peptide by all-atom molecular dynamics (MD) simulations. In 1 µs long simulations with bound PAD, the side chain of E593 of the PAD peptide formed strong and persistent salt-bridge interactions with a basic cluster on CK1 formed by Arg127, Lys154 and Lys171 (Fig. 5a, 5b, Supplementary Fig. 5a and 5b). By stabilizing the crystallographic position of the peptide, these interactions contribute to the slow phosphorylation rate of S591. This finding is consistent with a mutational analysis of the PAD peptide showing a ~two-fold acceleration for the E593A mutant (Fig. 3e, Supplementary Fig. 3c, 3d and Supplementary Table 1). Mutating the residues of the basic cluster on CK1 individually to glutamate and measuring the phosphorylation kinetics revealed that mutation of Lys154 and Lys171 resulted in a 1.5-fold faster kinetics of the third and fourth CK1 phosphorylation reactions (Supplementary Fig. 5c and 5d). These results confirmed the contribution of the interaction between E593 and the kinase to the slow phosphorylation rate of S591.

**Fig. 5.**
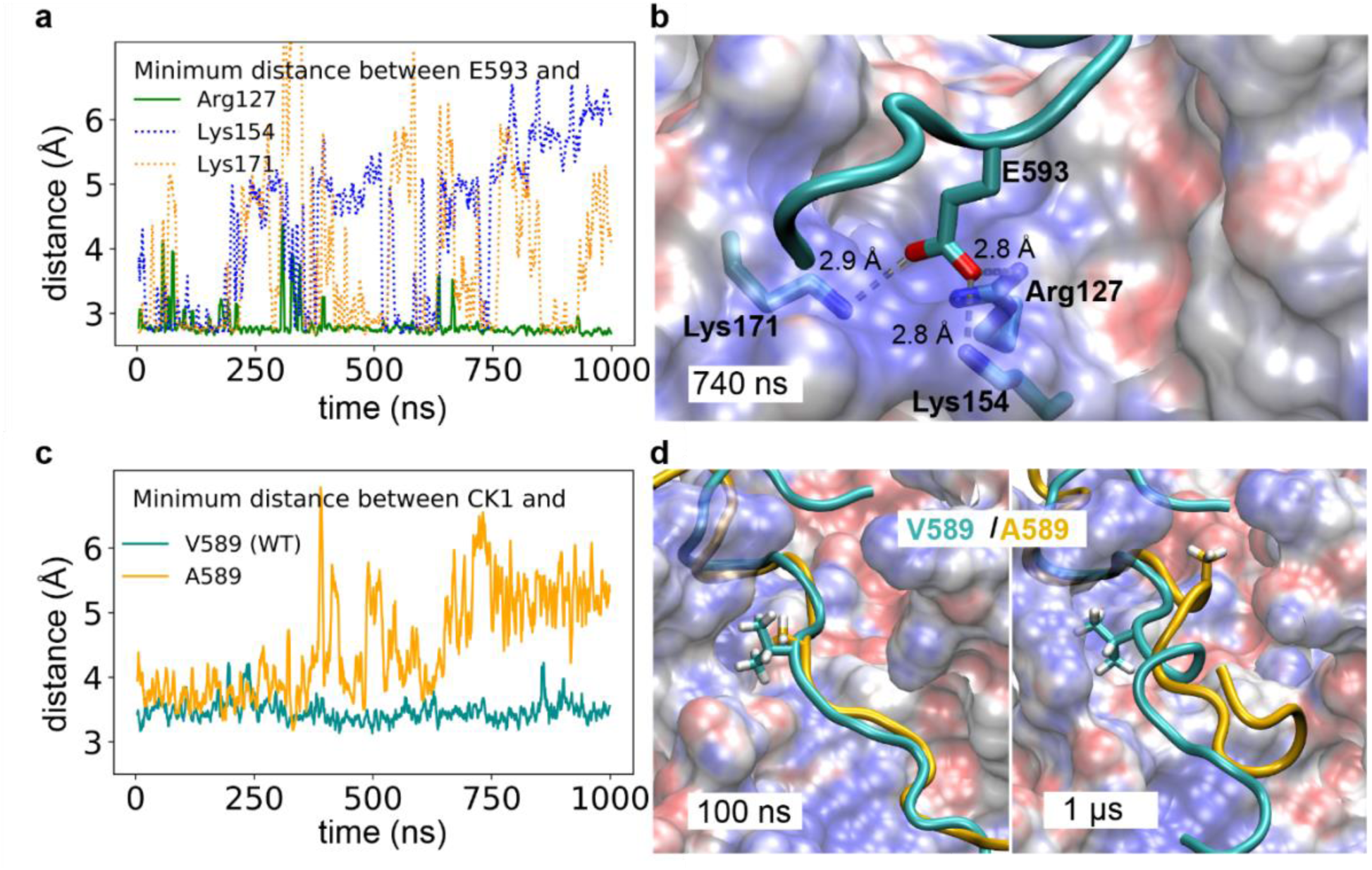
MD Simulations indicate that E593 and V589 of the p63 peptide are important for interaction with CK1. **a**, E593 is pinned down by salt bridges with CK1. Minimum heavy-atom distances of E593(p63) to Arg127(CK1) and Lys154(CK1) as function of time. E593 forms a salt bridge with Lys154 (distance < 3 Å), and binds transiently to Lys154. **b**, Representative snapshot at 740ns, zooming in on the C-terminal region of the p63 peptide. CK1 is shown as transparent electrostatic surface (blue/red for positive/negative charge) and the p63 peptide is represented as cyan cartoon. The residues E593, Arg127 and Lys154 are highlighted. The minimum distances between E593 and the basic residues are indicated. **c**, Mutation of V589 to A589 enhances the flexibility of the p63 peptide. Minimum heavy-atom distances of CK1 to V589 and A589 in WT and mutant p63 peptides as function of time. V589 remains adhered to the CK1 protein surface, whereas A589 gets solvated and moves away from the CK1 protein surface. **d**, Snapshots at 100 ns and 1 µs, zooming in on the C-terminal region of the p63 peptide. CK1 structures were superimposed. CK1 from the wild type simulation is represented as in **b** and the p63 peptide is shown in cartoon representation (cyan: WT; yellow: V589A) with V589 and A589 highlighted. Shown are results for the longer p63 construct. See Supplementary Fig. 5 for the capped and shortened p63 construct.

MD simulations also highlighted the stabilizing interactions of V589 on the PAD peptide with CK1. We compared two 1 µs long simulations of the WT and V589A PAD peptide. The V589A mutation weakened the interactions with a hydrophobic patch on CK1 and increased the flexibility of the peptide in this region (Fig. 5c and 5d). This finding is consistent with our mutational analysis, which showed that a large hydrophobic residue in the i+1-position retains the peptide and thereby slows down phosphorylation at the third CK1 site (Fig. 3e).

The presence of PAD-1P, PAD-2P and PAD-3P in the crystal structures suggests that all PAD peptides must have high binding affinities for CK1δ. However, we hypothesized that they may not exhibit the same binding and/or substrate potencies. To investigate this hypothesis, we measured k_cat_, K_m_ and V_max_ values for the phosphorylation reaction starting with single, double or triple phosphorylated peptides by measuring ATP consumption. Since the experiment was conducted with the wild type peptide sequence and not serine to alanine mutants the obtained values represent average values for all remaining phosphorylation events. These measurements clearly demonstrate that the affinity of the peptides increases and the reaction velocity decreases with increasing phosphorylation level (Supplementary Fig. 5e).

### The phosphorylation kinetics of the third CK1 site sets the threshold level for activation and oocyte death

Beyond the mechanistic description of the activation process, the question of the biological purpose of the observed kinetics remains. Analysis of the phosphorylation kinetics both of the isolated peptide as well as the full-length protein showed that phosphorylation of the third CK1 site starts only after the first two sites are completely phosphorylated (Fig. 2b and 2e). The first two sites act as a buffer creating a preloaded state and the third one acts as the trigger. Delaying the irreversible spring-loaded activation might allow the cell to survive in case the DNA damage does not surpass a certain threshold. This could be achieved by phosphatases that act during the delay period and remove the phosphorylation sites that constitute the buffer. One additional safety mechanism to control the fate of the oocyte is the degradation of activated TAp63α. Analysis of the total TAp63α concentration in oocytes has revealed that it decreases following the activation to the active tetrameric state^4,9^. A fast degradation in combination with a slow activation could also prevent apoptosis in case the DNA damage level is minor. To investigate the potential effects of dephosphorylation, degradation and delay of activation we simulated the entire process mathematically according to the kinetic model shown in Fig. 6a, Supplementary Fig. 6a and 6b. For simplicity, we used only one kinetic constant, k_1_, to describe the phosphorylation by CHK2 and the first two CK1 phosphorylation events. The kinetic constants k_2_ and k_4_ describe the dephosphorylation and degradation processes while k_3_ represents the critical third CK1 phosphorylation event that converts the inactive, dimeric protein [B] into the open, active conformation [C]. We solved the indicated system of coupled differential equations by numerical integration for different values of the kinetic constants. In our *in vitro* phosphorylation experiments the value of k_1_ was constant depending only on the concentration of CK1. However, *in vivo*, k_1_ will change with time as it depends for example on the amount of DNA damage, the activation kinetics of CHK2 as well as DNA repair processes. To better address this situation, we used different time-shifted curves as a description of the time dependence of the k_1_ value (Supplementary Fig. 6c, 6e and 6g). This resulted in a shift in the time response curve of [C], reflecting the experimental data of p63 tetramerization in oocytes but had otherwise little effect on [C] (Supplementary Fig. 6d, 6f and 6h). Changing the value for k_2_ also showed relatively little effect on the formation of the activated tetramer [C] (Fig. 6b). In contrast, k_4_ has a strong effect with high values keeping the concentration of activated tetramer low (Fig. 6c).

**Fig. 6.**
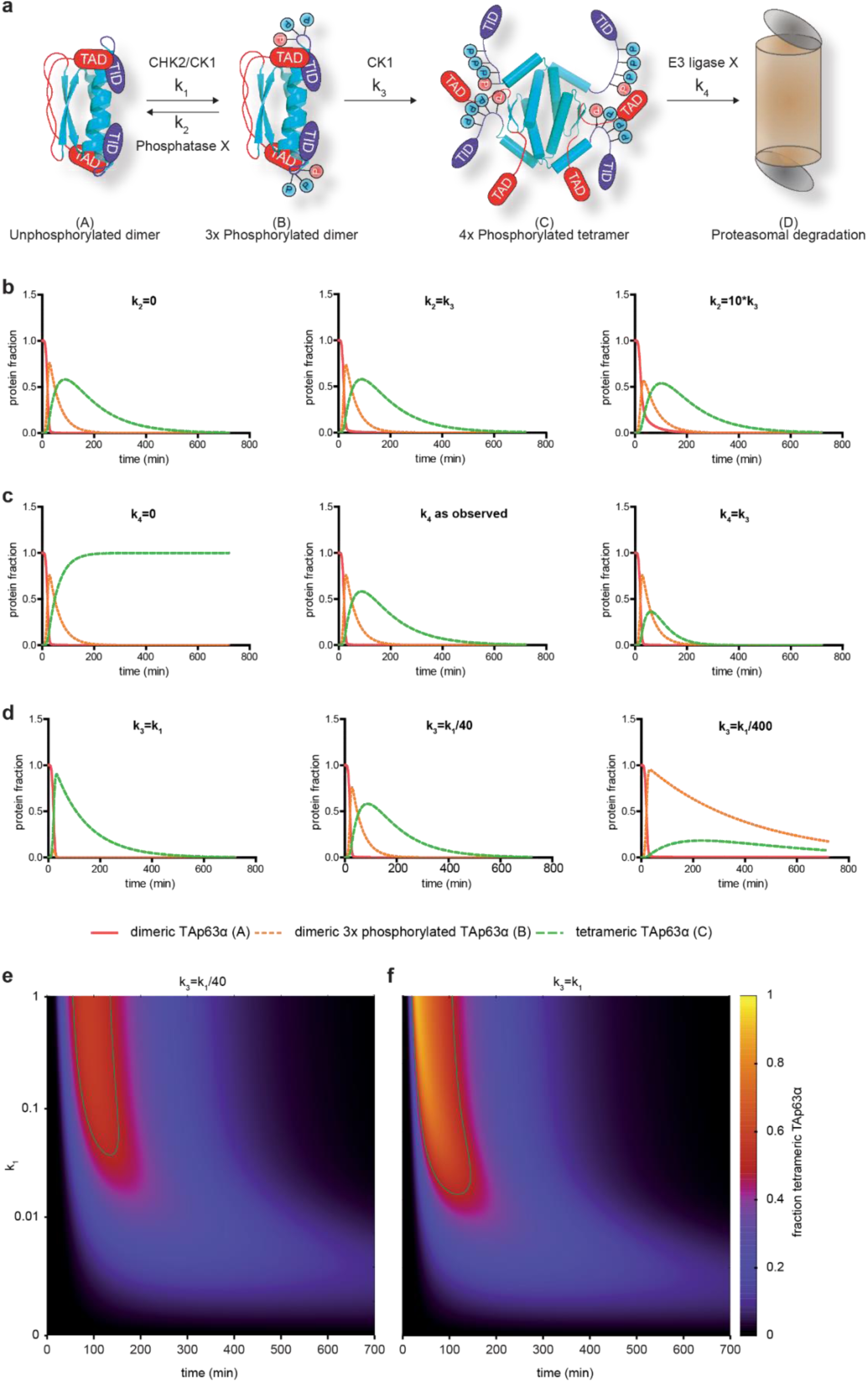
Simulation of the effect of different kinetic constants on the concentration of tetrameric active TAp63α [C]. **a**, Model of the phosphorylation dependent activation of TAp63α. Phosphorylation by CHK2 and the first 2 CK1 sites are combined into one kinetic constant k_1_. Dephosphorylation and degradation are described by k_2_ and k_4_, respectively. The decisive third phosphorylation by CK1 is represented by kinetic constant k_3_. **b**, Curves representing the concentration of non-phosphorylated, dimeric TAp63α [A] (red), triple phosphorylated, dimeric TAp63α [B] (yellow) and quadruple phosphorylated, tetrameric TAp63α [C] (green) were calculated according to the model in **a** for three different values of k_2_, indicated above each diagram. **c**, same as in **b** but varying the value of k_4_. **d**, same as in **b** and **c** with variation of k_3_. **e**, Two-dimensional plot of time (x-axis) vs k_1_ (y-axis). Variations of k_1_ are shown on a logarithmic scale. The concentration of [C] is represented on the z-axis with different colors according to the values shown at the right. The green line represents the contour level of a [C]-value of 0.5 which was set arbitrarily as the level of activation that would induce apoptosis (‘zone of death’). **f**, Same as in **e** but with a 40-times faster k_3_ level showing that the ‘zone of death’ is larger. See also Supplementary Fig. 6.

These simulations predict that phosphatases probably play a less important role than E3 ligases in regulating of TAp63α mediated oocytes death. To address the decisive question of the influence of the k_3_ kinetic constant we selected values for k_4_ based on the observed decay of the tetrameric p63 level in oocytes following irradiation (k_4_=0.008 min^−1^; Supplementary Fig. 1a) and calculated the time course of the concentration of activated tetramer for low and high values of k_3_. These calculations show a strong influence of the level of [C] on the value of k_3_ (Fig. 6d). With k_3_ values approaching k_1_, meaning that no lag phase in phosphorylation exists, a fast rise of [C] to high concentrations results. Slower k_3_ values dampen the response of the system and delay it, enabling the degradation process to effectively remove activated tetramer. To better characterize the effect of the phosphorylation lag phase we calculated the time response curve for [C] in dependence of varying the maximal value of k_1_. This calculation mimics an almost instant and persistent level of DNA damage as seen following γ-irradiation. Low k_1_ values represent low DNA damage while large k_1_ values represent strong DNA damage. The result is a three-dimensional plot with time and k_1_ values as x and y axis and the level of [C] on the z axis. Fig. 6e shows a two-dimensional representation with contour lines for the value of [C]. Fig. 6f shows the same calculation only with k_3_ set to a higher level equal to the maximal value of k_1_. Setting a [C]-value of 0.5 arbitrarily as the level that has to be reached to induce apoptosis demonstrates that this ‘zone of death’ is smaller for the calculations with the slow k_3_ value. Our simulation results suggest that the delayed phosphorylation of S591 sets the threshold for the DNA damage level that is necessary to induce apoptosis.

## DISCUSSION

Cell division and cell death require yes/no answers without the possibility for an intermediate response. Consequently, cells have evolved molecular systems that translate ‘analog’ input signals into a ‘digital’ output signal. Many theoretical and experimental studies have shown different ways how cells build switch-like systems, often using multisite phosphorylation events as a signal integrator of the activity of two or more kinases. Although activation of TAp63α requires two different kinases, this integrator model does not seem to be the mode of activation, because CK1 kinases are thought to be constitutively active but sequentially dependent on a priming kinase such as CHK2. In this situation, activation of TAp63α would only depend on the activation of CHK2. This model, however, is only valid if CK1 used a strictly processive mode of phosphorylation with all individual kinetic constants in the same range. Our results show, that both conditions are not true and that CK1 uses a distributive mode of phosphorylation in combination with very different kinetic constants to regulate TAp63α’s activation. Importantly, it is the third phosphorylation event that is the slowest and that constitutes the decisive phosphorylation for the formation of the open, tetrameric state. The transition to the open tetrameric state is irreversible and all phosphate groups can be removed in this state without affecting its oligomerization status^9^. Phosphorylation of the third CK1 site, therefore, constitutes the ‘point of no return’. Mechanistically the slow kinetics seems to be the result of three effects. 1) S592 and E593 interact with a basic / polar cluster of the kinase and probably keep the peptide in the position as seen in the crystal structures. 2) Hydrophobic residues such as valine are preferred in the +1-position relative to the phosphorylation site due to the presence of a small hydrophobic patch next to the active site. Shifting S592 into this position would not allow a stable interaction. 3) The triple phosphorylated peptide (pS582, pS585, pS588) has a relatively high affinity to the kinase in a product type state. The crystal structure of the triple phosphorylated peptide with the phosphorylated S588 still pointing into the active site suggested product inhibition might play a role in slowing down phosphorylation of S591. This interpretation is supported by a study showing that temperature compensation of the circadian clock of which CK1 is a central component is at least partially based on a shift from high substrate affinity to higher product affinity of the kinase^26^, showing that product inhibition is an important aspect of regulation of CK1 activity.

Sequence alignments of the p63 PAD peptides revealed that not only the phosphorylation sites are highly conserved from fish to mammals^13^ but also the valine-glycine sequence following the second CK1 site and the serine – glutamate sequence C-terminal to the third site, suggesting that activation of TAp63α follows a universal, evolutionary conserved mechanism with similar activation kinetics in all these species.

An important question of multisite phosphorylation events is whether kinases use a processive or distributive mechanism. Examples exist for both types as well as a combination of both. Src kinase and SRPK1 have been shown to modify up to fifteen tyrosine and eight serine residues in a processive manner, respectively^27,28^ while the dual specificity kinase MEK uses a fully distributive mechanism^23^. One example for a combined mechanism is the yeast cyclin-dependent kinase Pho85 that modifies five residues in the transcription factor Pho4 using both modes^29^. Our experiments have demonstrated that CK1 uses most likely a distributive mechanism of phosphorylation. The first two phosphorylation events are too fast to analyze in detail but more likely represent a distributive mechanism in combination with a fast rebinding. The third site, however, follows definitely a distributive and not a processive mode. Effectively, the first two CK1 phosphorylation sites act as a buffer that gets filled with a relatively fast kinetics. Only when these two sites are phosphorylated do the third (and fourth) sites become modified, resulting in a sigmoidal time response curve. Importantly, the general behavior of the individual phosphorylation events is not restricted to the relatively artificial phosphorylation of isolated peptides but can also be seen in the kinetic analysis of the same sequence within the full-length TAp63α protein. Originally, we conducted these experiments with the full-length protein because we wanted to investigate potential steric hindrance effects. Surprisingly, phosphorylation of the third CK1 site is not slower, but faster within the full-length protein. This result might be a consequence of the dimeric state of TAp63α, presenting two binding sites for CK1 per molecule (in addition to potentially many other non-specific interaction sites) that might keep the kinase in close proximity to TAp63α, increasing the local concentration and in the end accelerating phosphorylation.

The model that we use to simulate the effect of the kinetic constants is of course simplified. Activated TAp63α will initiate transcription of many other genes including p63 itself and probably E3 ligases which will influence the cellular concentration and localization. Despite these simplifications our results show that the difference in phosphorylation kinetics sets the threshold for p63 activation using a ‘doorbell like’ mechanism. This threshold level has been established during evolution to suppress oocyte loss due to random activation of TAp63α or low levels of DNA damage caused for example by background radioactivity or normal levels of reactive oxygen species in oocytes. Our model predicts that mutations in the PAD sequence of TAp63α can change this threshold level. A single nucleotide exchange within the codon of S592, AGT, to ATT would mutate this serine to an isoleucine, which would change the phosphorylation kinetics and, therefore, the threshold, leading to TAp63α activation at lower damage levels. Two recent screens have shown, that mutations in TAp63α can lead to POI. In both screens large rearrangements in the domain structure have created activated p63 forms^30,31^. A S592I mutation would have much milder effects but could also contribute to a premature depletion of the primary oocyte reserve.

## ACKNOWLEDGMENTS

The authors would like to thank Ina Theofel, Sarah Young and Eric Chih-Chao Liang for their review of and input into this manuscript. The research was funded by the DFG (DO 545/8-1), the Centre for Biomolecular Magnetic Resonance (BMRZ), and the Cluster of Excellence Frankfurt (Macromolecular Complexes). M.T. was supported by a Fellowship from the fund of the German Chemical Industry. L.S. and G.H. were supported by the Max Planck Society. F.P., K.H. and E.H.K.S. thank the EU Horizon2020 project LSFM4LIFE (grant # 668350-2), the ZonMw-BMBF joint sponsored project ‘The Onconoid Hub’ (grant # 114027003) for funding. The Structural Genomics Consortium is a registered charity (number 1097737) that receives funds from the Canadian Institutes for Health Research, the Canadian Foundation for Innovation, Genome Canada through the Ontario Genomics Institute, GlaxoSmithKline, Karolinska Institute, the Knut and Alice Wallenberg Foundation, the Ontario Innovation Trust, the Ontario Ministry for Research and Innovation, Merck & Co., Inc., the Novartis Research Foundation, the Swedish Agency for Innovation Systems, the Swedish Foundation for Strategic Research, and the Wellcome Trust.

## AUTHOR CONTRIBUTIONS

J.G. and M.T. performed NMR and kinetic assays *in vitro* and in ovaries. A.C. crystallized and solved the structures of the kinase-peptide complexes. K.H. and F.P. performed microscopy experiments and quantitative semi-automated segmentation, L.S. carried out MD simulations. F.L. performed NMR experiments, N.G. measured tetramerization kinetics, F.F., E.H. and J.M. expressed and purified proteins, M.S. measured phosphorylation kinetics, R.L. solved differential equations. M.T., G.H., E.H.K.S., S.K. and V.D. designed experiments and analyzed data. J.G., M.T., A.C. and V.D. wrote the manuscript.

## MATERIALS & METHODS

### Ovary Culture

Animal care and handling were performed according to the guidelines set by the World Health Organization (Geneva, Switzerland). Eight-day-old (P8) female CD-1 mice were purchased from Charles River Laboratories. Ovaries were harvested, transferred to sterile 96-well plates with 50 µl α-MEM (+L-Glu, Gibco) supplemented with 10% FBS (Gibco), 1x penicillin/ streptomycin (Gibco), 0.2 mg/ml Na-pyruvate (Gibco), 2 mg/ml *N*-acetyl-l-cysteine (Sigma) and ITS liquid media supplement (100x) (Sigma) cultured at 37°C with 5% CO_2_ overnight^32^. The final concentrations of the kinase inhibitors targeting ATM (KU55399, Selleckchem), CHK2 (BML-277, Merck) and CK1 (PF 670462, Sigma Aldrich) were 25 µM. *Ex vivo* ovaries were γ-irradiated with 0.50 Gy on a rotating turntable in a ^137^Cs irradiator with a dose of 2.387 Gy/min. The protocol for harvesting mouse ovaries was approved by the Tierschutzbeauftragte of the Goethe University Frankfurt/Main.

### Western blotting and blue native PAGE

Western blotting and blue native PAGE (BN-PAGE) were performed as described previously^13,33^. The following antibodies were used for detection: anti-p63α (D2K8K XP, Cell Signaling), anti-cleaved-PARP (D6X6X, Cell Signaling), anti-VASA (DDX-4) (ab13840, Abcam).

### Whole Ovary staining, optical clearing and light sheet-based fluorescence microscopy (LSFM)

Cultured ovaries were treated as indicated and the staining was performed as described in Tuppi et al.^13^. In brief, the ovaries were harvested and fixed in 4% paraformaldehyde (PFA) in PBS overnight at 4°C. The ovaries were permeabilized using 0.3% Triton X-100 in PBS for 30 minutes at room temperature. All further steps were performed in a 96-well flat bottom plate (Greiner) shaking at 450rpm. The ovaries were blocked for two hours using Blocking Buffer (0.3% Triton X-100, 0.05% Tween-20, 0.1% BSA (heat shock fraction, Sigma) and 10% Donkey Serum in PBS). Then the ovaries were incubated overnight at 37°C in a humidified incubator with the first antibody, an anti-Germ cell-specific antigen (GCNA1) (ab82527, Abcam), 1:200 and anti-cleaved PARP (D6X6X, Cell Signaling) and 1 µg/ml DAPI, which was diluted in the Blocking Buffer. This was followed by three 20 minutes PBS washes and a subsequent four hours incubation of the secondary antibody, donkey anti-rabbit Alexa 488 (A-21206, Thermo Fisher) and donkey anti-rat Alexa 568 (ab1754775, Abcam), in a 1:200 dilution in Blocking Buffer containing 1 µg/ml DAPI at 37°C in a humidified incubator in the dark. Protected from light, the ovaries were then washed three times for 20 minutes with PBS and kept at 4°C in PBS. The ovaries were cleared by washing them four times in CUBIC 2^34^ (50% w/v Sucrose, 25% w/v Urea,10% w/v 2,2’,2’’-nitrilotriethanol) with a refraction index of 1.49 and overnight incubation in CUBIC 2 inside a fluorinated ethylene propylene (FEP) foil capillary (Patent: US 20150211981 A1). The capillaries were mounted on stainless steel holders and images were acquired with a custom-built monolithic digital-scanned light sheet-based fluorescence microscope (mDSLM)^35^. The microscope was equipped with an Epiplan-Neofluar 2.5x/0.06 illumination objective and a N-Achroplan 10x/0.3 detection objective (Carl Zeiss) and a Clara camera (ANDOR technology, Ireland). Z-spacing of 2.58 µm. Laser and filter set: 488 nm laser, 525/50 bandpass filter. All raw image stacks were pre-processed in FIJI (ImageJ version 1.51d, Java version 1.6.0_24). Specifically, the raw image stacks were cropped to the region of interest and scaled by a factor of two. To obtain a homogeneous intensity distribution in the individual images of the z-stacks, the stacks were resliced by a factor of four. The background intensity was subtracted using the FIJI function Subtract Background with a ball radius of 50 pixels and the contrast was enhanced applying the FIJI function Enhance Contrast (saturation=0.35). GCNA and c-PARP positive cells were segmented with the FIJI function 3D Object Counter. Objects with a size of min.=750 and max.=222218880 voxels and an intensity threshold of 1,000 were segmented. The total amount of c-PARP and GCNA positive cells per ovary were extracted and the ratio of GCNA/c-PARP per ovary was calculated. Three replicates per time point were analyzed.

### TAp63α expression, purification and phosphorylation

Human TAp63α (aa10-616, TEV site inserted at aa570 and V599M) codon-optimized for expression in *E. coli* was subcloned into the pET16b vector. The protein, bearing a C-terminal His_6_-tag was expressed in BL-21(DE3)-R3-Rosetta (SGC Oxford) for 16 h at 18°C in M9 minimal Media, containing 1 g/l ^15^NH4Cl as nitrogen source. Additionally, the media was supplemented with 100 µM ZnCl_2_ to ensure correct folding of the zinc finger containing DNA binding domain. Cells were lysed in IMAC buffer A (50 mM Tris pH 8.0, 400 mM NaCl, 20 mM β-mercaptoethanol, 5% glycerol, 10 µM ZnCl_2_) and purified using a standard step gradient (300 mM imidazol) immobilized metal affinity chromatography (IMAC) protocol (Ni-Sepharose Fast Flow, GE Healthcare). Afterwards, the protein was further purified by size exclusion chromatography (SEC) (HiLoad 16/600 Superdex 200 pg, GE Healthcare) in 50 mM Tris pH 8.0, 150 mM NaCl, 5% glycerol, 0.5 mM TCEP. Purified TAp63α was incubated for 30 min at 30°C with pre-activated MK2Δ1-41, 10 mM ATP and 10 mM MgCl_2_. Purification and activation of MK2Δ1-41 was performed as described previously ^13^. Subsequently, TAp63α was separated from MK2 by SEC (HiLoad 16/600 Superdex 200 pg, GE Healthcare). 2.5 mg of the pre-phosphorylated TAp63α (pTAp63α) was then incubated with purified CK1δ (molar ratio 1:1000) with 10 mM ATP and 10 mM MgCl_2_ at 25°C for the indicated time. Purification and activation of CK1δ was performed as described previously^13^. To stop the reaction PF670462 (10 µM final concentration) and EDTA pH 8.0 (12 mM final concentration) were added. Afterwards, MBP-TEV were added in a 1:1 molar ratio and incubated overnight at 4°C. The cleavage was stopped by adding solid urea to it to a final concentration of 6 M. The denaturated mixture was concentrated and purified by SEC (Superdex 75, 10/300 GL, GE Healthcare) in 50 mM Tris pH 8.0, 150 mM NaCl, 6 M Urea. The resulting PAD-TID peptide was acidified with 32% HCl to a final concentration of 0.5 M and cleaved by a final concentration of 125 mM CNBr for 48 h at room temperature in the dark. The resulting product was concentrated for 30 min in a SpeedVac. Afterwards, MES pH 6.3 was added to a final concentration of 500 mM to adjust the pH. To remove the TID the reaction was purified by a reverse IMAC, concentrated and buffer exchanged (kinase NMR buffer, see below) by ultrafiltration (Amicon Ultra-0.5 ml, 3 kDa MWCO, Merck) for NMR analysis.

### Peptide Expression

Peptides were expressed with a protease cleavable N-terminal GFP-His_6_-tag. Isotopically labeled expression was performed in M9 for 16 h at 22°C under the induction with 500 µM IPTG. Initial purification was done analogous to full-length TAp63α. After IMAC purification, the expression tag and the peptide were split by incubation with 3C protease overnight at 4°C. On the next day the peptide and GFP were subject to concentration via ultrafiltration (Amicon Ultra 10 kDa MWCO, Merck) and removal of the GFP and 3C protease. The flow through of the filter was subject to another round of concentration over a 3 kDa cutoff filter. Afterwards the peptides were subject to SEC in kinase NMR buffer, see below (Superdex 75 10/300 GL, GE Healthcare).

### Peptide MK2 pre-phosphorylation

Peptides were S582 -phosphorylated with MK2 kinase at 25°C at a molar ratio of 1:100 kinase to peptide and an ATP concentration of 10 mM. Phosphorylation was monitored by recording a series of 2D NMR spectra. MK2 was subsequently removed from the reaction mix by another round of SEC (Superdex 75, 10/300 GL, GE Healthcare).

### NMR Spectroscopy

Samples for NMR experiments employed kinase buffer (50 mM Bis-Tris pH 6.5, 50 mM NaCl, 10 mM MgCl_2_). Additionally, all samples contained 1× protease inhibitor cocktail (cOmplete EDTA free, Roche) and 1× phosphatase inhibitor cocktail (PhosSTOP, Roche). Experiments were performed at a sample temperature of 298 K. Assignments of PAD mutants at different phosphorylation states were performed using a combinatorial triple-selective labeling approach, as detailed in^36^ or with the help of a constant-time HNCACB.

Phosphorylation kinetics were recorded using sample volumes of 200 µl placed in 3-mm capillaries and final peptide concentrations of 250 µM. The general sample composition of a kinetic sample was as follows:

|  |  |
| --- | --- |
| <sup>15</sup>N-labeled peptide | to 250 μM |
| protease inhibitor | 1x |
| phosphatase inhibitor | 1x |
| ATP | to 10 mM |
| CK1 kinase (3 μM stock) | 8.3 μl |
| buffer | to 200 μl |

A series of HSQC-like [^15^N, ^1^H] correlation spectra were recorded throughout the kinetic reaction with NMR instruments operating at ^1^H frequencies ranging from 600 to 950 MHz. To accurately reflect the extremely fast initial phosphorylation reactions of S582 and S585 the total measurement time of each spectrum was limited to a maximum of 120 s. This was achieved by using a gradient-selected BEST-TROSY pulse sequence^37^ and either by limiting the measurement to one scan per FID or by reducing the interscan delay to 100 ms, depending on the field strength the sample was measured at. Non-uniform sampling was not employed as the large quantity of ATP within the sample leads to non-recoverable t1-noise like artefacts, overlaying the signals.

Prior to addition of kinase a reference spectrum was acquired. Kinase was added manually, and a series of spectra were recorded over the time course of ~12 h. Peak intensities of relevant peaks were quantified for every spectrum automatically. The fraction of phosphorylation in a given amino acid was calculated by dividing the relevant peak intensity/intensities by the sum of all peak intensities of a given amino acid.

YAP1 phosphorylation kinetics were measured based on ^13^C detected 2D (H)NCO experiments to resolve peak overlap in the [^15^N, ^1^H]-HSQC spectrum. Two-dimensional ^13^C detected N-CO correlation spectra were obtained using a BEST-(H)NCO pulse sequence essentially as described by Gil et al.^38^. Nitrogen chemical shifts were acquired in a semi-constant time manner using States-TPPI quadrature detection. Homonuclear ^13^CO-^13^Cα decoupling was achieved by the IPAP approach^39^. To allow for very short interscan delays without causing sample - and RF coil heating ^15^N decoupling during acquisition was not applied in the current implementation. Acquisition was immediately started after the 9.4-ms IPAP delay (= (2^1^*J*_CαCO_)^−1^), leading to a slightly shorter pulse sequence duration. Refocusing of the active ^1^*J*_CON_ coupling takes place during the fixed ^13^CO-^13^Cα IPAP delay and proceeds during acquisition. As a consequence, line shapes along the directly detected ^13^CO dimension are a superposition of non-resolved in-phase ^1^*J*_CON_ doublets, phased to absorption, and dispersive antiphase ^1^*J*_CON_ doublets. It should be noted that higher resolution in the carbonyl dimension could be obtained using virtual ^13^CO-^15^N decoupling as originally proposed^38^, but this would entail doubling the experimental time because twice the number of FIDs would have to be recorded for each time domain data point. Spectra were recorded on a Bruker AVIII 800 MHz spectrometer equipped with a cryogenic ^1^H/^13^C/^15^N triple resonance TXO probe optimized for ^13^C detection. Acquisition times were 60 ms for ^13^C and 98.7 ms for ^15^N with spectral widths of 20 ppm in both dimensions. Four FIDs were acquired for each time domain data point in the indirect dimension (two for ^15^N quadrature detection and two for ^13^CO-^13^Cα IPAP) and two scans were accumulated for each FID. Using a relaxation delay of 0.1 s and non-uniform sampling (37.5 % sparse) the total experimental time for each spectrum was 2 min. Phosphorylated Ser/Thr residues were identified on the basis of ^*3*^*J*_CαP_ couplings using a ^31^P-edited intra-HNCA experiment as previously proposed^40^. To enable a direct comparison with the N-CO correlation spectra acquired to monitor the phosphorylation reaction, a carbonyl evolution time (t1) was introduced here, resulting in the 3D [^15^N, ^1^H]-BEST-TROSY-COintraHN(CAP) pulse sequence shown in Supplementary Fig. 3b. The projection along the ^1^H dimension of the final 3D spectrum provides the desired ^15^N_i_-^13^CO_i-1_ cross peaks of phosphorylated residues *i*. The spectrum was acquired with a cryogenic ^1^H/^31^P/^13^C/^15^N quadruple resonance QCI probe on a Bruker AVIIIHD 700 MHz spectrometer. Spectral widths were adjusted to 8, 10 and 9.6 ppm, respectively, along the ^13^C, ^15^N, and ^1^H dimensions, where the ^1^H carrier was placed on the water frequency. Acquisition times were 53.8 ms (^13^C, 76 complex points), 115.5 ms (^15^N, 82 complex points), and 76.1 ms (^1^H, 512 complex points). Non-uniform sampling was employed to record a total of 2181 hypercomplex points (35% of the full *t*_*1*_/*t*_*2*_ grid). The spectrum was acquired within 24 h using a recycle delay of 0.25 s and 16 scans/FID.

### NMR kinetic analysis

The phosphorylation kinetics was analyzed according to a sequential model:

[A] -> [B] -> [C] -> [D] with [A] representing the CHK2-phosphorylated PAD peptide and [B], [C] and [D] representing successive phosphorylated species. The functions for fitting the experimental data were:

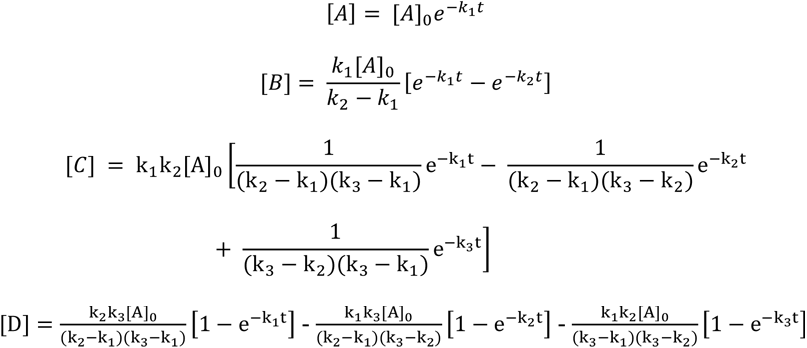

Fitting and visualization were performed with the software package Prism 6.0 (GraphPad).

### Crystal structure determination

The kinase domain of CK1δ (aa. 1-294) was subcloned into pNIC28-Bsa4, and the recombinant protein containing an N-terminal His_6_ tag was expressed in *E. coli*, cultured in TB and induced with 0.5 mM IPTG at 18°C overnight. Cells were harvested and lysed by sonication, and the protein was purified by IMAC. The histidine tag was cleaved by TEV protease treatment, and the cleaved protein was purified further by reverse IMAC and SEC. The pure protein in 20 mM Tris, pH 7.5, 200 mM NaCl and 0.5 mM TCEP was concentrated to 6-8 mg/ml, and was mixed with either ADP or AMPPCP (Sigma) at 2 mM and MgCl_2_ at 4 mM. Crystallization was performed using sitting drop vapor diffusion method at 4°C using the condition containing 10-20% PEG 3350, 0.1-0.2 M sodium sulfate and 0.1 M citrate, pH 4.6-5.9. Viable crystals were soaked with the PAD peptides at 6-10 mM overnight in the mother liquor containing 20% ethylene glycol before flash-cooled in liquid nitrogen. Diffraction data were collected at Swiss Light Source, X06SA or DESY, P13, and were processed and scaled with XDS^41^ and subsequently scaled using aimless^42^, respectively. The CK1δ structures in complexes with the peptides were solved by molecular replacement using Phaser^43^ and the published coordinates of CK1δ ^44^. Manual model rebuilding alternated with structure refinement was performed in COOT^45^ and REFMAC^46^, respectively. Geometry correctness of the final structures were checked using Molprobity^47^. Data collection and refinement statistics are summarized in Supplementary Table 2.

Peptides used for crystallization in complex with CK1δ:

PAD-1P = YTP(pS)SASTVSVGSSET (MW = 1639.6, e = 1490)

PAD-2P = YTP(pS)SA(pS)TVSVGSSET (MW = 1719.6, e = 1490)

PAD-3P = (pS)SA(pS)TV(pS)VGSSY (MW 1371.16, e=1490)

### Mathematical model

The system of differential equations describing the model in Fig. 6 was solved numerically using the Runge-Kutta method implemented in the software package SageMath 8.7. The script used is shown below.

For Fig. 6 b-d the following script was used:

w,x,y,z,t=var(‘w x y z t’)

a=1/(1+(30/(t+0.000001))^4) # k_1_

b=0.008 # k_2_

c=1/40 # k_3_

d=0.008 # k_4_

P=desolve_system_rk4([-a*w+b*x,a*w-(b+c)*x,c*x-d*y,d*y],[w,x,y,z],ics=[0,1,0,0,0], ivar=t,end_points=720,step=1)

Q1=[[i,j] for i,j,k,l,m in P]

Q2=[[i,k] for i,j,k,l,m in P]

Q3=[[i,l] for i,j,k,l,m in P]

print table(columns=[Q1,Q2,Q3])

For Fig. 6 e,f the following script was used:

w,x,y,z,t=var(‘w x y z t’)

o = 0

v = 0

while o <= 1000:

v=(1000^(o/1000)-1)/999 a=(1/(1+(30/(t+0.000001))^4))*v # k_1_

b=0.008 # k_2_

c=1/40 # k_3_

d=0.008 # k_4_

P=desolve_system_rk4([-a*w+b*x,a*w-(b+c)*x,c*x-d*y,d*y],[w,x,y,z],ics=[0,1,0,0,0], ivar=t,end_points=720,step=1)

Q3=[[o,i,l] for i,j,k,l,m in P]

print Q3

o = o + 1

The different functions for k_1_ were as follows:

sigmoidal curve with initial delay: a=1/(1+(30/(t+0.000001))^4) constant k1: a=1

hyperbolic k_1_: a=(exp(1)-exp(1/t))/(exp(1)-1)

Gaussian curve k_1_: a=exp(-((t-40)/20)^2)

Curves were plotted using the software package Prism 6.0 (GraphPad). Two-dimensional plots were created with Gnuplot (http://www.gnuplot.info/).

### ADP-Glo

k_M_, V_max_ and k_cat_ values for given kinase substrates were determined using the ADP-Glo Kit system (Promega). In all cases reactions were carried out in 384 well plates at room temperature for 30 min with a total kinase concentration of 50 nM and 1 mM ultra-pure ATP. To limit the concentration error due to the extremely low assay volume (5 µl) peptide dilution series as well as ATP and CK1δ addition were performed by an Echo 550 (Labcyte) ultrasonic liquid handler. Phosphorescence was detected with a Spark plate reader (Tecan).

### Molecular Dynamics Simulations

Initial simulation models were built according to the X-ray crystal structure p63-PAD-3P, with bound ADP replaced by ATP. All crystallographic water molecules and ions within 10 Å of the protein were retained. ATP and a complexed Mg^2+^ ion were added using the Protein Data Bank entry 1CSN as template, by superimposing the protein backbones and aligning the nitrogen atoms of ATP with the crystallographic ADP. Missing side chains were added using the software Modeller^48^. The triple phosphorylated peptide PAD-3P was elongated to TPpSSApSTVpSVGSSETRGER with charged termini as used in the kinetic measurements by means of VMD Molefracture^49^ and Modeller (S592 and E593 coordinates from p63-PAD-2P). Using these tools, we also constructed a complex of CK1 with a shortened peptide ACE-TPpSSApSTVpSVGSSETRG-NME capped with N-terminal acetyl and C-terminal methylamino capping groups. This peptide mimics the full-length protein, because E597 and R598 may not be accessible to CK1. In a third setup, the point mutation V589A was introduced into the longer peptide using Modeller. In all three setups, Asp128 was protonated. All other residues were simulated in their physiological protonation state. All three MD simulations were carried out with Gromacs 2018^50^ using the AMBER99SB*-ILDN-q force field^51–54^, the water model TIP3P^55^ and the Schwierz ion force field^56^ for Mg^2+^ and NaCl at 150 mM concentration. Force field parameters for ATP and phosphoserine were taken from Meagher et al.^57^ and Homeyer et al.^58^, respectively. The system was energy minimized followed by five equilibration steps with successively decreasing position restraints on heavy atoms, first in an NVT ensemble (0.25 ns) and then in an NPT ensemble (4 x 0.5 ns) using a Berendsen thermostat and barostat^59^. The three production runs of 1 µs each were run at a temperature of 310 K in an NPT ensemble using a Nosé-Hoover thermostat^60^. The pressure was maintained at 1 bar with a Parrinello-Rahman barostat^61^. The minimum heavy-atom distances between E593 and Arg127, between E593 and Lys154, as well as the minimum heavy-atom distance between V589 or A589 (sidechain) and the protein CK1 were monitored at 1 ns intervals using the gmx mindist tool. The raw distance data were processed using Moving Average Smoothing with a window size of 5. All structural figures were made using VMD^49^.

### Data accessibility

All data are fully available upon request.

---

**Supplementary Fig. 1.**
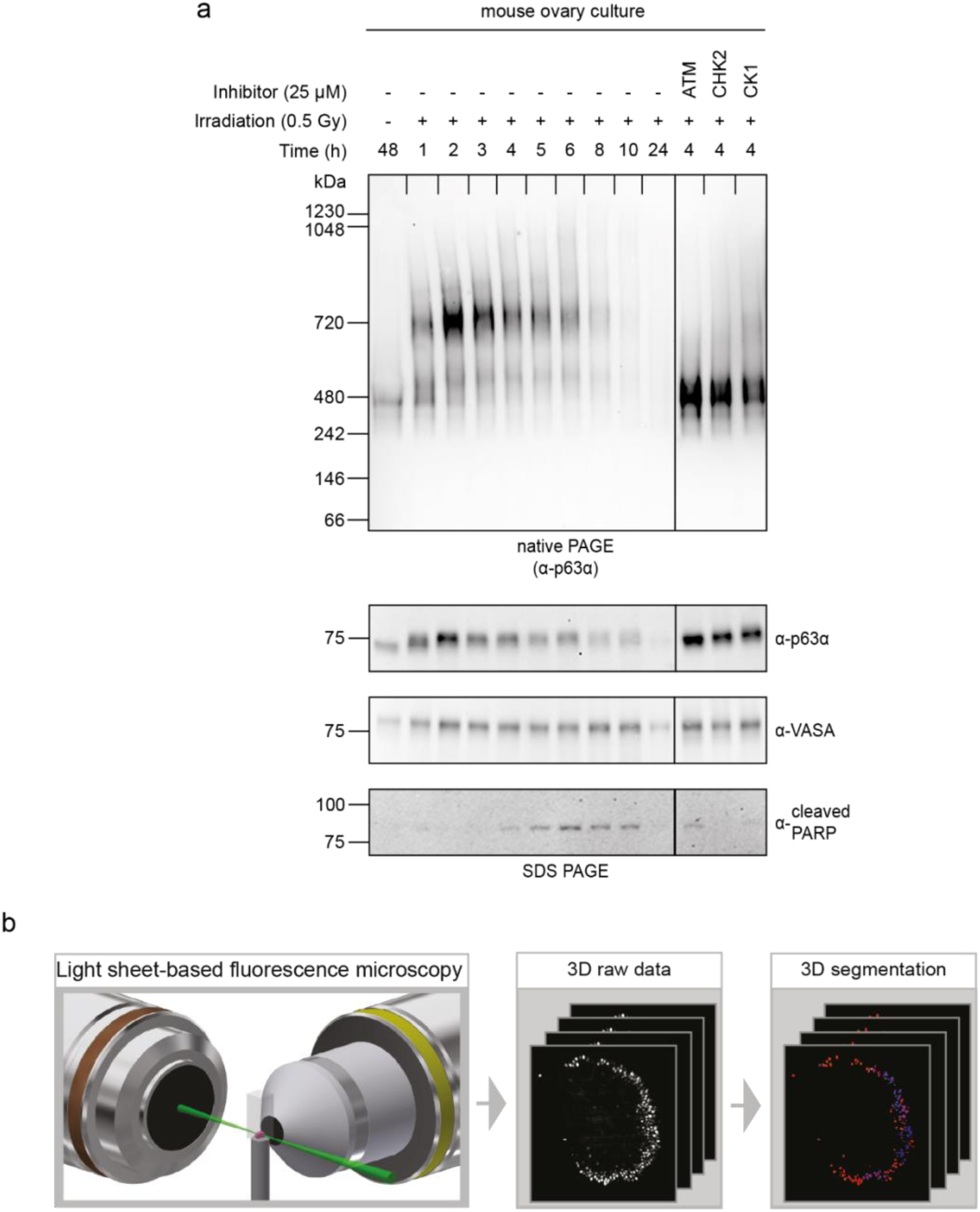
**a**, Whole mouse ovaries were *ex vivo* γ-irradiated with 0.5 Gy with or without indicated kinase inhibitors or left untreated. The oligomeric state was analyzed as a function of time by BN-PAGE and the phosphorylation dependent mobility shift was analyzed by SDS-PAGE. Apoptosis was monitored by detecting cleaved PARP and primordial oocytes level were detected using α-VASA antibody. **b**, Workflow of our developed light sheet-based whole mount 3D ovary staining. Ovaries were stained and imaged using FEP foil holders and light sheet-based fluorescence microscopy. The raw data were segmented and semi-automatically processed resulting in a quantitatively analyzed evaluation in whole ovaries.

**Supplementary Fig. 2.**
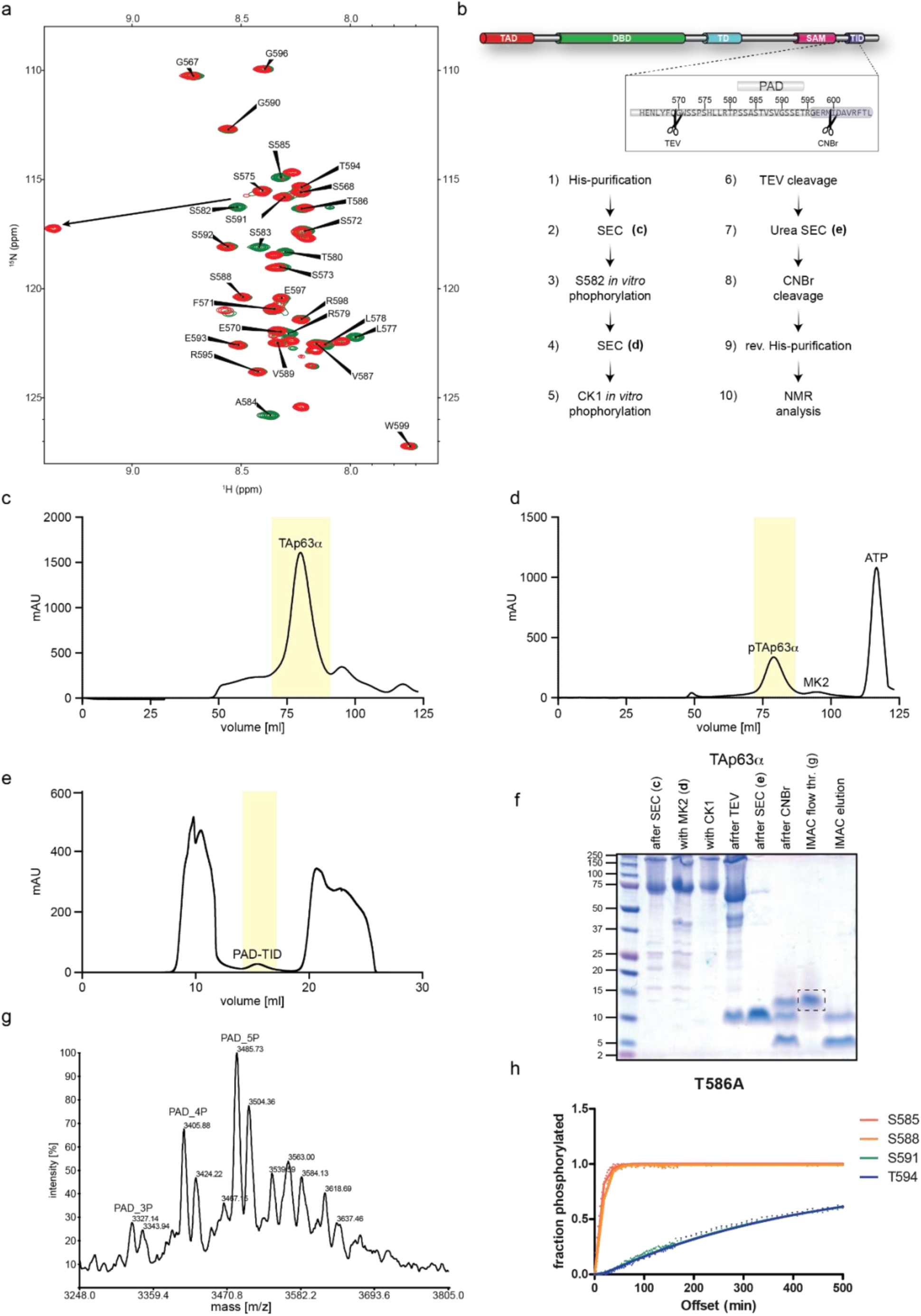
**a**, Overlay of HSQS spectra of unphosphorylated (green) and MK2 phosphorylated PAD peptide (red). A large chemical shift change in S582 can be observed as the result of phosphorylation. **b**-**d**, General purification strategy of phosphorylated PAD peptides from full length TAp63α. The protein was expressed in *E. coli* and initially purified via IMAC, followed by SEC (Superdex 200 16/600) (**c**). Dimeric fractions (highlighted yellow) were collected and subject to MK2 phosphorylation. This was followed by another round of SEC (Superdex 200 16/600) to remove MK2 kinase (**d**). **e**, Denaturing SEC (Superdex 75 10/300 GL) of cleaved TAp63α. The PAD-TID peptide is marked in yellow. **f**, SDS-page summarizing the purification of the PAD peptide from full-length protein. The dashed box corresponds to the peptide analyzed by mass spectrometry in **g. g**, ESI spectrum of the PAD peptide isolated from TAp63α. Several different phosphorylation states of the peptide can be observed. Generally, a set of two peaks, with a mass difference of 18 Da, can be observed for each phosphorylation state. This most likely is the result of partial hydrolysis of the homoserine-lactone created by CNBr cleavage of the PAD-TID sequence. **h**, T586A mutant of the PAD peptide, showing the same distinct difference between S588 and S591 phosphorylation kinetics as the wild type sequence.

**Supplementary Fig. 3.**
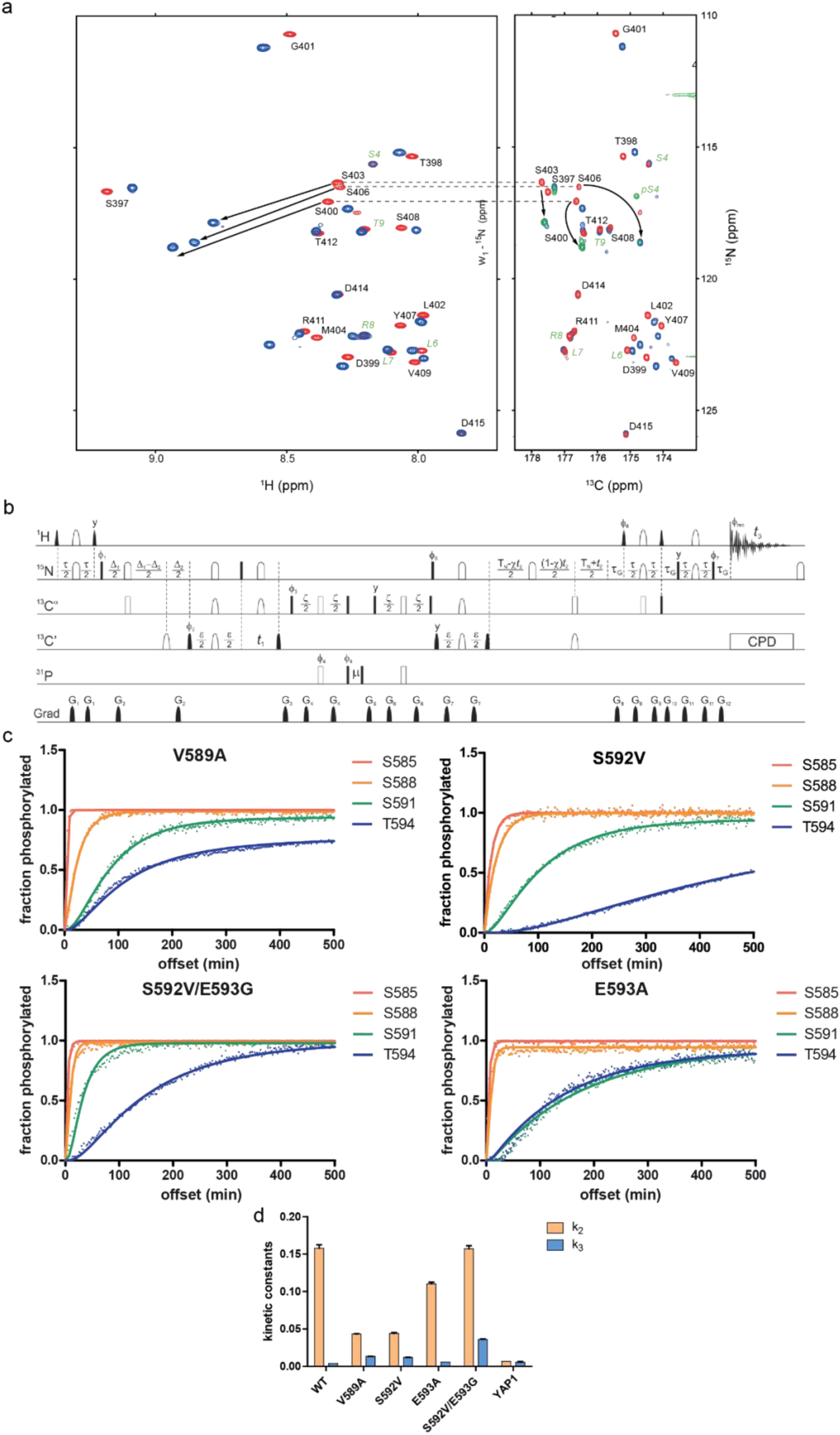
**a**, Comparison of 2D HSQC (left) and 2D ^13^C-detected NCO (right) planes. In the mono-phosphorylated state (red) S403 and S406 are almost completely degenerated in the HSQC while in the (H)NCO they can be completely resolved. In the fully phosphorylated case (blue) all peaks can be resolved in the HSQC as well as in the (H)NCO. To determine which residues are indeed phosphorylated a 2D projection (NCO plane) of the experiment described below was used. The resulting projection is shown as green overlay in the right panel. **b**, Experimental scheme of the 3D [^15^N, ^1^H]-BEST-TROSY-COintraHN(CAP) experiment. Radiofrequency (RF) pulses with flip angles of 90° and 180° are represented by filled and open symbols, respectively. Offsets for ^1^H, ^15^N, ^13^C^α, 13^C’, and ^31^P are 8.7, 118.5, 56.7, 176.7, and 3.5 ppm, respectively. The initial two proton 90° pulses have a PC9 shape, the third and fourth use time-reversed E-BURP-2 and regular E-BURP-2 shapes, respectively, and a RE-BURP shape is employed for ^1^H 180° pulses. Bandwidths of all proton pulses are adjusted to 4 ppm. Rectangular ^15^N 90° pulses are applied with the highest available power, while 180° flip angles are achieved with broadband inversion pulses (BIPs) of 0.7 ms duration, except for the last one before acquisition, which uses of RE-BURP shape with a bandwidth of 40 ppm. Carbonyl-selective pulses (both 90 and 180°) of 0.103 ms duration (at 700 MHz spectrometer frequency) have an amplitude envelope corresponding to the center lobe of a sin(x)/x function. Rectangular pulses on α-carbons are applied off-resonance using phase modulation with an RF field of v/15^1/2^ for 90° flip angle and v/3^1/2^ for 180° flip angle, where v is the difference between ^13^C^α^ and ^13^C’ offsets in Hz. The simultaneous 180° pulses on α- and carbonyl carbons in the center of ε periods are implemented as double-band RE-BURPs with inversion/refocusing bands of ca. 16 ppm (C’ region) and ca. 22 ppm (C^α^ region), separated by 120 ppm. Delay durations are as follows: τ = 5.4 ms, Δ_1_ = 38 ms, Δ_2_ = 33 ms, ε = 9 ms, ζ = 55 ms, µ = 0.003 ms, T_N_ = 33 ms, and τ_G_ = 0.39 ms. The *t*_2_ evolution time is implemented in a semi-constant time manner where X = T_N_/*t*_2,max_. The default RF pulse phase is x. Phase cycling: Φ _1_ = 16(y), 16(−y); Φ _2_ = 4(y), 4(−y); Φ _3_ = x, −x; Φ _4_ = 2(x), 2(−x); Φ_5_ = 8(x), 8(−x); Φ _rec_ = X, 2(−X), X, −X, 2(X), −X, where X = x, 2(−x), x. Pulsed field gradients along the z-axis have a sine-bell shape and durations of 0.3 ms (G1-G7 and G9), 0.175 ms (G8, G10, and G12), and 1.0 ms (G11), respectively. Peak amplitudes are G_1_: −10 % (percentage of maximum available gradient strength); G_2_: −9 %; G_3_: −15 %; G_4_: −12 %; G_5_: 12%; G_6_: −14 %; G_7_: 8 %; G_9_: −28 %; G_11_: 65 %. Coherence selection is achieved by gradients G_8_, G_10_, and G_12_ that are applied with amplitudes (−70 %, 30 %, 19.87 %) and (−80 %, 20 %, 30.13 %) to record N- and P-type transients, respectively. For each *t*_2_ increment both types are collected alternately by changing pulse phases Φ_6_ from y to –y and Φ_7_ from x to –x. The two FIDs are stored separately and then added and subtracted to form the real and imaginary parts of a complex data point with a 90° zero-order phase shift being added to one of the components. Phase Φ_5_ is inverted along with the receiver reference phase in every other increment to shift axial peaks to the edge of the spectrum in the ^15^N dimension. Quadrature in the ^13^C dimension was achieved by applying States-TPPI to Φ_2_. **c**, Phosphorylation kinetics of selected PAD mutants showing a reduction in the kinetic difference between S588 and S591. **d**, Measured kinetic constants for the phosphorylation of S588 (k_2_) and S591 (k_3_) in wild type and different mutant PAD peptides as well as the constants for the second and third phosphorylation event in the YAP1 peptide.

**Supplementary Fig. 4.**
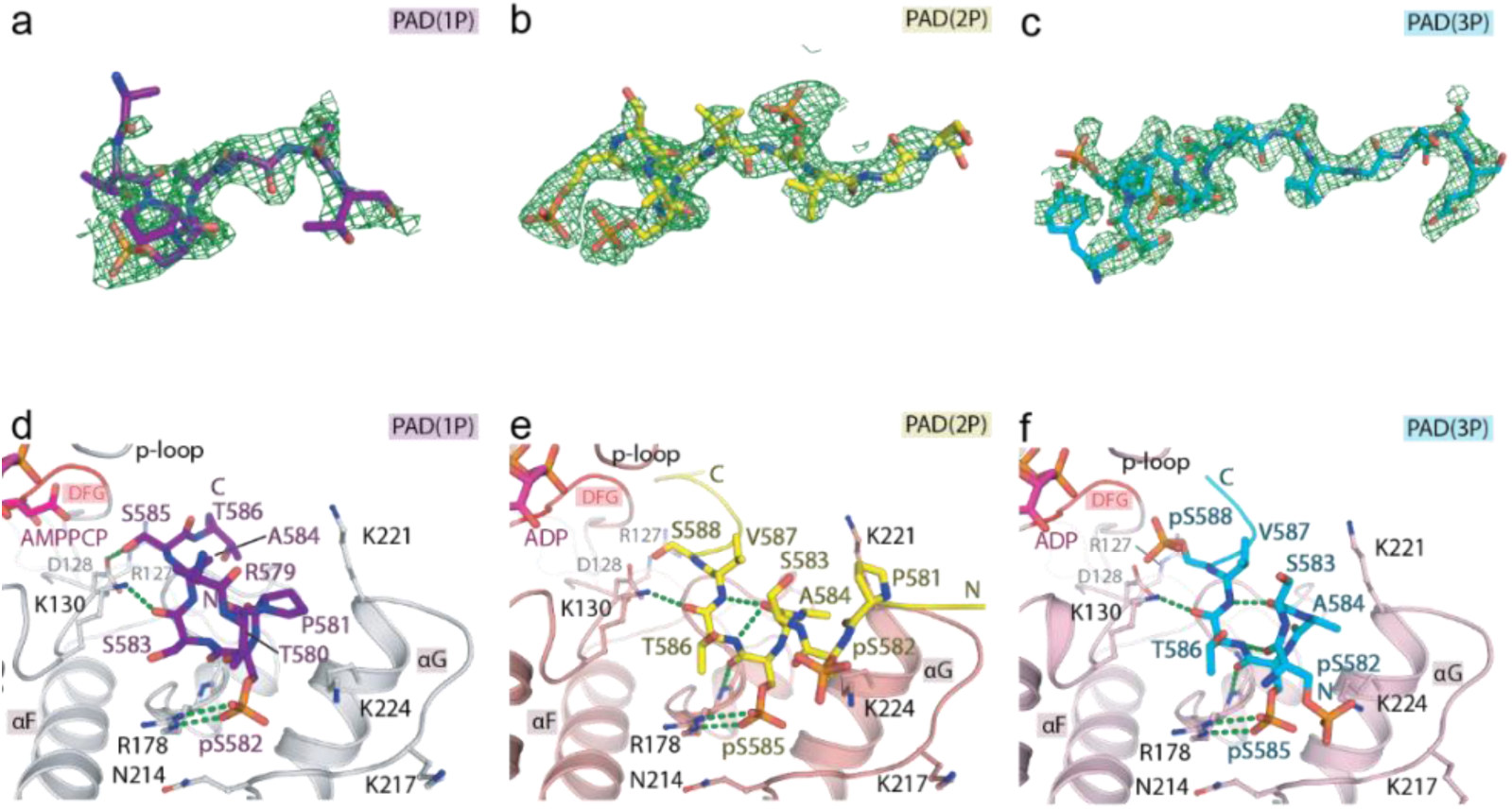
**a-c**, Binding of the PAD peptides within CK1δ in the crystal structures. |Fo|-|Fc| omitted electron density map contoured at three-sigma for the bound PAD peptides. **d**-**f**, Detailed interactions at the N termini of the PAD peptides within the kinase.

**Supplementary Fig. 5.**
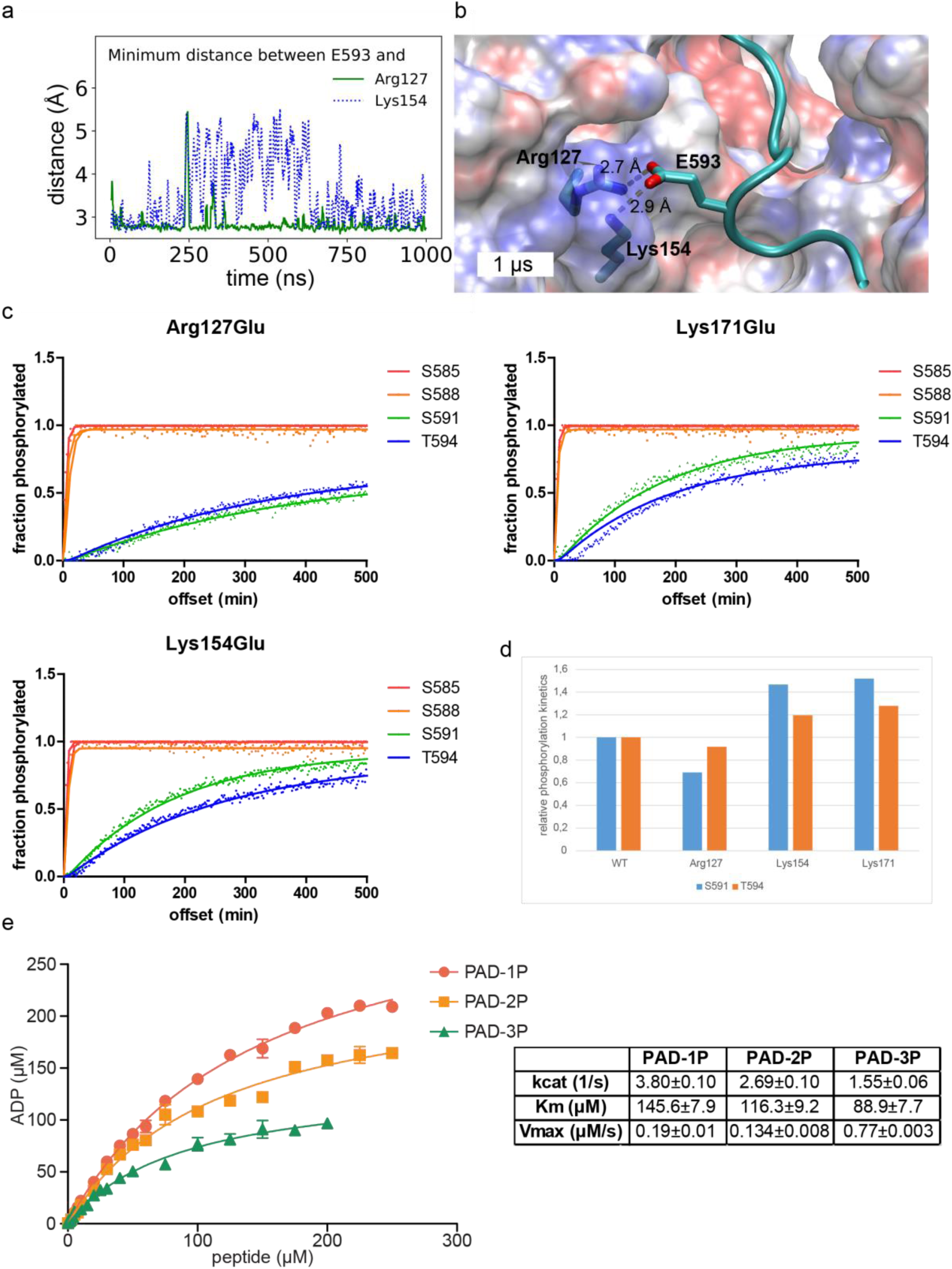
**a**, MD simulation of CK1 in complex with a shorter PAD-3P peptide (ACE-TPpSSApSTVpSVGSSETRG-NME) with N-terminal acetyl and C-terminal methylamino capping groups showing similar results as the longer peptide (Fig. 5a and 5b). **b**, Snapshot at 1 µs, zooming in on the C-terminal region of the shorter p63 peptide. CK1 is shown as transparent electrostatic surface (blue/red for positive/negative charge) and the p63 peptide is represented as cyan cartoon. The residues E593, Arg127 and Lys154 are highlighted. The minimum distances between E593 and the basic residues are indicated. **c**, Phosphorylation kinetics of selected CK1δ mutants showing a reduction in the kinetic difference between S588 and S591 compared to the wild type kinase. **d**, Relative phosphorylation kinetics of the mutants described in **a**, normalized on wild type kinase. Mutation of Lys154Glu shows the strongest difference in phosphorylation kinetics for T594 while the difference is largest for S591 in case of a Lys171Glu mutation. **e**, Determination of K_M_, v_max_ and k_cat_ for mono-, di- and tri-phosphorylated PAD peptides. The assay was performed in triplicates, the indicated error represents the standard deviation.

**Supplementary Fig. 6.**
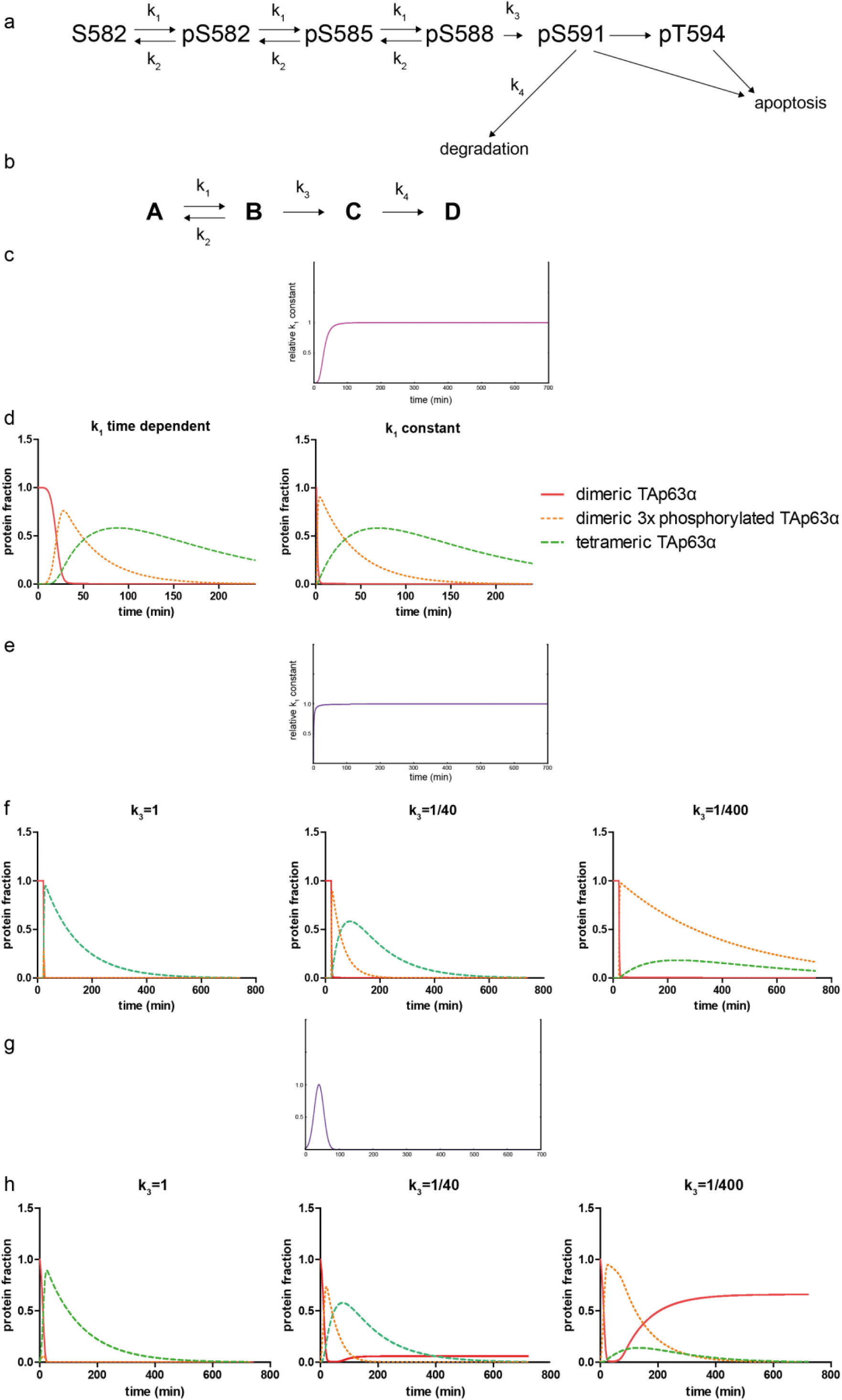
**a**, Simplified kinetic model of the activation of TAp63α. Upon DNA damage TAp63α is modified by CHK2 kinase and subsequently by CK1. For this model it is assumed that the initial CHK2 phosphorylation events as well as the first two CK1 modifications are fast and can therefore be summarized into a single kinetic constant k_1_. The hypothetical activity of phosphatases counteracting the phosphorylation at this point is described as k_2_. The decisive phosphorylation of S591 is described by k_3_ and the competitive reaction to apoptosis, namely degradation of the tetrameric protein, is described as k_4_.**b**, Further simplified diagram of the kinetic model described in **a. c**, Plot of the time dependent function used to model the initial DNA damage response (k_1_). **d**, Comparison of time dependent k_1_ as used in the model and a constant k_1_. **e**, Plot of an alternative, hyperbolic k_1_ used for simulated kinetics shown in **f. f**, Influence on different k_3_ kinetic constants on the formation of tetrameric protein under a hyperbolic k_1_. The start of the kinetics has been artificially set to 20 min to account for the experimentally observed lag after γ-irradiation of ovaries. **g**, Plot of an alternative, Gaussian k_1_ used for simulated kinetics shown in panel **f. h**, Influence on different k_3_ kinetic constants on the formation of tetrameric protein under a Gaussian k_1_

**Supplementary Table 1.** Kinetic constants derived from the NMR based phosphorylation experiments.

| peptide | k<sub>1</sub> | k<sub>2</sub> | k<sub>3</sub> | k<sub>4</sub> |  | k<sub>2</sub>/WT k<sub>2</sub> | k<sub>3</sub>/WT k<sub>3</sub> | k<sub>2</sub>/k<sub>3</sub> |
| --- | --- | --- | --- | --- | --- | --- | --- | --- |
| WT | 0.2688 | 0.1581 | 0.0038 | 0.0033 |  | 1.00 | 1.00 | 41.99 |
| V589A | 0.3002 | 0.0434 | 0.0134 | 0.0099 |  | 0.27 | 3.55 | 3.24 |
| S592V | 0.0797 | 0.0441 | 0.0120 | 0.0020 |  | 0.28 | 3.18 | 3.69 |
| E593A | 0.2147 | 0.1103 | 0.0057 | 0.0067 |  | 0.70 | 1.52 | 19.26 |
| S592V/E593G | 0.2639 | 0.1573 | 0.0359 | 0.0719 |  | 0.99 | 9.55 | 4.38 |
| YAP | 0.0072 | 0.0066 | 0.0054 | - |  | 0.04 | 1.43 | 1.23 |

**Supplementary Table 2.**
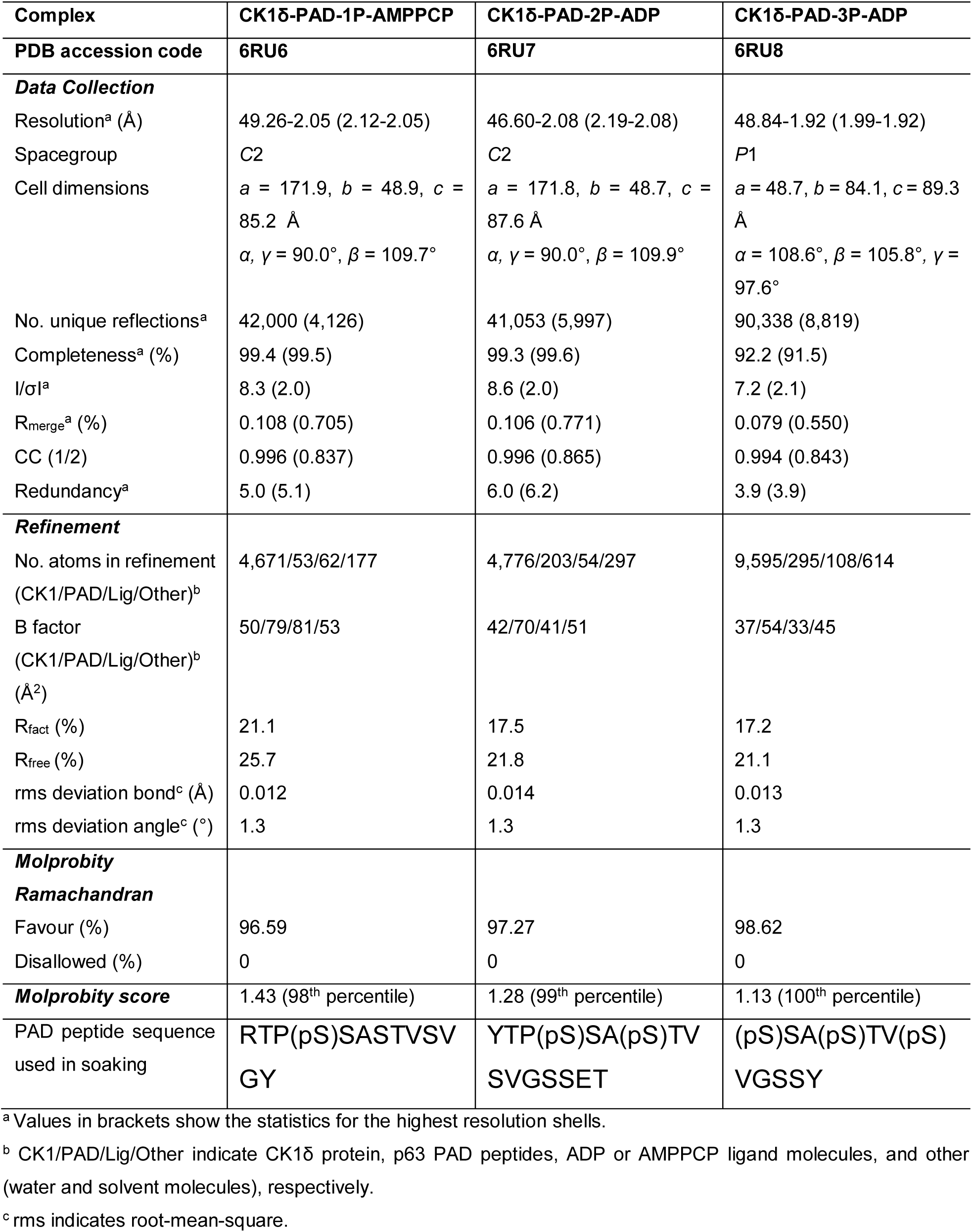
Data collection and refinement statistics.

| Complex | CK1δ-PAD-1P-AMPPCP | CK1δ-PAD-2P-ADP | CK1δ-PAD-3P-ADP |
| --- | --- | --- | --- |
| PDB accession code | 6RU6 | 6RU7 | 6RU8 |
| **Data Collection** |  |  |  |
| Resolution<sup>a</sup> (Å) | 49.26-2.05 (2.12-2.05) | 46.60-2.08 (2.19-2.08) | 48.84-1.92 (1.99-1.92) |
| Spacegroup | C2 | C2 | P1 |
| Cell dimensions | $a = 171.9, b = 48.9, c = 85.2$ Å<br>$\alpha, \gamma = 90.0^\circ, \beta = 109.7^\circ$ | $a = 171.8, b = 48.7, c = 87.6$ Å<br>$\alpha, \gamma = 90.0^\circ, \beta = 109.9^\circ$ | $a = 48.7, b = 84.1, c = 89.3$ Å<br>$\alpha = 108.6^\circ, \beta = 105.8^\circ, \gamma = 97.6^\circ$ |
| No. unique reflections<sup>a</sup> | 42,000 (4,126) | 41,053 (5,997) | 90,338 (8,819) |
| Completeness<sup>a</sup> (%) | 99.4 (99.5) | 99.3 (99.6) | 92.2 (91.5) |
| I/σI<sup>a</sup> | 8.3 (2.0) | 8.6 (2.0) | 7.2 (2.1) |
| R<sub>merge</sub><sup>a</sup> (%) | 0.108 (0.705) | 0.106 (0.771) | 0.079 (0.550) |
| CC (1/2) | 0.996 (0.837) | 0.996 (0.865) | 0.994 (0.843) |
| Redundancy<sup>a</sup> | 5.0 (5.1) | 6.0 (6.2) | 3.9 (3.9) |
| **Refinement** |  |  |  |
| No. atoms in refinement (CK1/PAD/Lig/Other)<sup>b</sup> | 4,671/53/62/177 | 4,776/203/54/297 | 9,595/295/108/614 |
| B factor (CK1/PAD/Lig/Other)<sup>b</sup> (Å<sup>2</sup>) | 50/79/81/53 | 42/70/41/51 | 37/54/33/45 |
| R<sub>fact</sub> (%) | 21.1 | 17.5 | 17.2 |
| R<sub>free</sub> (%) | 25.7 | 21.8 | 21.1 |
| rms deviation bond<sup>c</sup> (Å) | 0.012 | 0.014 | 0.013 |
| rms deviation angle<sup>c</sup> (°) | 1.3 | 1.3 | 1.3 |
| **Molprobrity** |  |  |  |
| **Ramachandran** |  |  |  |
| Favour (%) | 96.59 | 97.27 | 98.62 |
| Disallowed (%) | 0 | 0 | 0 |
| **Molprobrity score** | 1.43 (98<sup>th</sup> percentile) | 1.28 (99<sup>th</sup> percentile) | 1.13 (100<sup>th</sup> percentile) |
| PAD peptide sequence used in soaking | RTP(pS)SASTVSV<br>GY | YTP(pS)SA(pS)TV<br>SVGSSET | (pS)SA(pS)TV(pS)<br>VGSSY |

<sup>a</sup> Values in brackets show the statistics for the highest resolution shells.
<sup>b</sup> CK1/PAD/Lig/Other indicate CK1δ protein, p63 PAD peptides, ADP or AMPPCP ligand molecules, and other (water and solvent molecules), respectively.
<sup>c</sup> rms indicates root-mean-square.

